# Characterization of non-coding variants associated with transcription factor binding through ATAC-seq-defined footprint QTLs in liver

**DOI:** 10.1101/2024.09.24.614730

**Authors:** Max F. Dudek, Brandon M. Wenz, Christopher D. Brown, Benjamin F. Voight, Laura Almasy, Struan F.A. Grant

## Abstract

Non-coding variants discovered by genome-wide association studies (GWAS) are enriched in regulatory elements harboring transcription factor (TF) binding motifs, strongly suggesting a connection between disease association and the disruption of cis-regulatory sequences. Occupancy of a TF inside a region of open chromatin can be detected in ATAC-seq where bound TFs block the transposase Tn5, leaving a pattern of relatively depleted Tn5 insertions known as a “footprint”. Here, we sought to identify variants associated with TF-binding, or “footprint quantitative trait loci” (fpQTLs) in ATAC-seq data generated from 170 human liver samples. We used computational tools to scan the ATAC-seq reads to quantify TF binding likelihood as “footprint scores” at variants derived from whole genome sequencing generated in the same samples. We tested for association between genotype and footprint score and observed 693 fpQTLs associated with footprint-inferred TF binding (FDR < 5%). Given that Tn5 insertion sites are measured with base-pair resolution, we show that fpQTLs can aid GWAS and QTL fine-mapping by precisely pinpointing TF activity within broad trait-associated loci where the underlying causal variant is unknown. Liver fpQTLs were strongly enriched across ChIP-seq peaks, liver expression QTLs (eQTLs), and liver-related GWAS loci, and their inferred effect on TF binding was concordant with their effect on underlying sequence motifs in 80% of cases. We conclude that fpQTLs can reveal causal GWAS variants, define the role of TF binding site disruption in disease and provide functional insights into non-coding variants, ultimately informing novel treatments for common diseases.

**Graphical Abstract:** 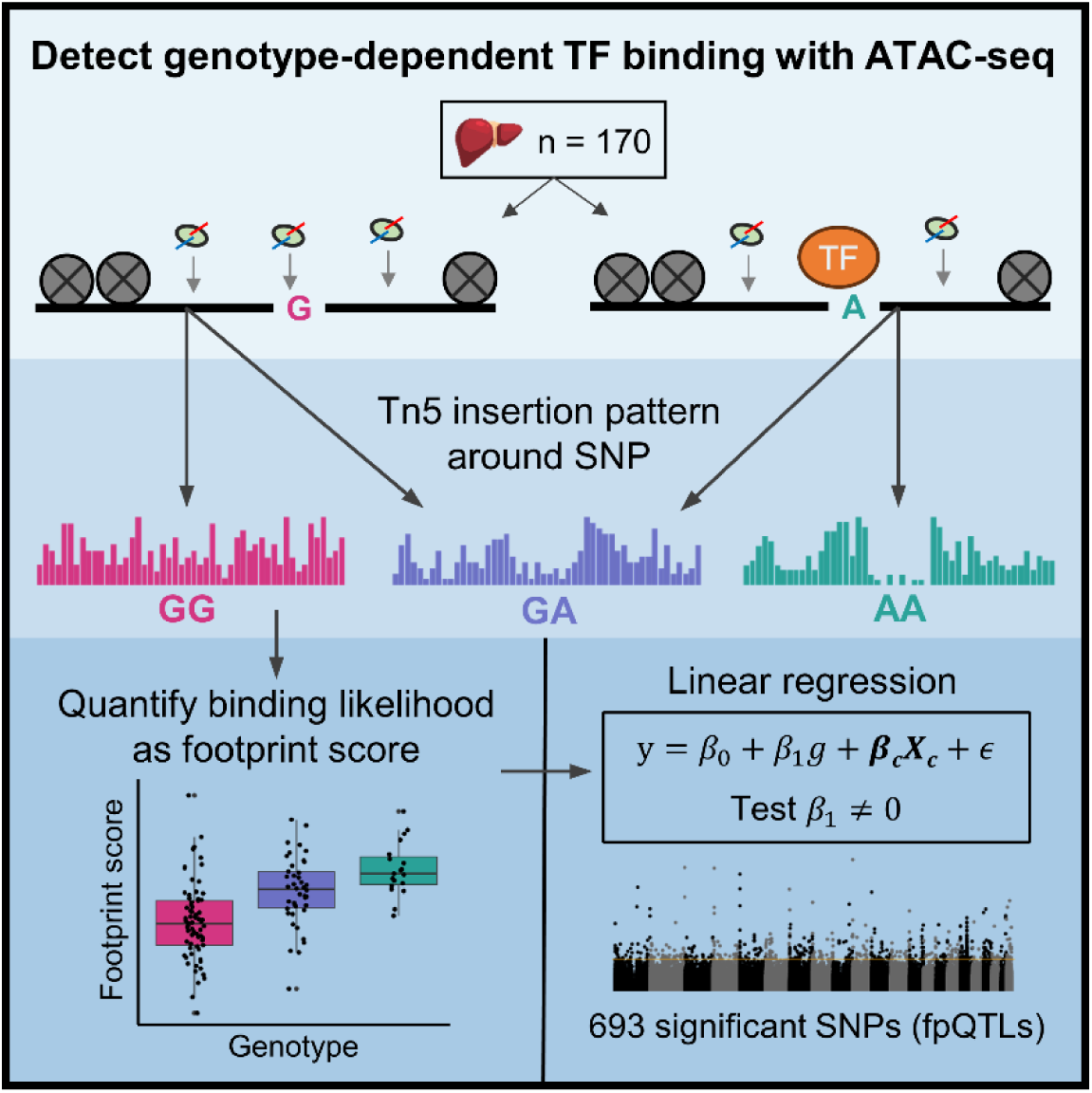

We leverage footprinting methods to infer transcription factor binding likelihood genome-wide across 170 liver ATAC-seq samples and implicate 693 SNPs with a genetic influence on binding. Unlike other comparable approaches, this analytical method is not limited in resolution by the constraints of linkage disequilibrium, and can prioritize likely causal variants at GWAS loci for subsequent experimental validation.

## Introduction

More than 90% of GWAS-implicated variants are located in non-coding genomic regions with uncharacterized effects on gene regulation^1–4^, limiting their utility in characterizing disease biology and implicating novel targets for treatment. Furthermore, given the structure of linkage disequilibrium (LD) across the genome, a variant with a true biological effect on some trait (i.e., a *causal* variant) will be correlated with nearby variants, making it challenging to distinguish which variants among them is causal^5–7^. A second challenge after association mapping is to determine the effector gene(s) regulated by a given causal variant through which the trait effect is conferred. Single nucleotide polymorphisms (SNPs) associated with gene expression, i.e. expression quantitative trait loci (eQTLs), as catalogued by the GTEx Consortium, are strongly linked with approximately 43% of disease-associated GWAS signals^8^, and on average only 11% of disease heritability is estimated to be explained by GTEx gene expression^9^. A recent study modeled that GWAS loci and eQTLs are systematically biased towards different types of cis-regulatory variants, suggesting that additional connections beyond those provided by eQTLs are needed to provide mechanistic insights into observed complex trait association signals^10^. Other variant-to-gene mapping approaches which use 3D chromatin architecture data from Hi-C or Capture-C require the causal variant to be nominated before implicating the effector gene^11–13^. Methods to experimentally validate the effects of putative causal SNPs on gene expression are expensive and time consuming, making the prioritization of candidate variants a key bottleneck in disease genomics.

Non-coding GWAS-implicated variants are concentrated in regulatory regions and near transcription factor (TF) binding motifs^1,14,15^, suggesting that the disruption of cis-regulatory sequence plays a mechanistic role in conferring disease risk. ATAC-seq, an experimental method traditionally used to measure chromatin accessibility, can also be used to detect TF binding. In this method, the transposase Tn5 inserts sequencing adapters into DNA, preferentially at genomic locations where chromatin is open^16^. However, bound TFs can partially block Tn5, leaving a pattern of relatively depleted Tn5 insertion sites known as a “footprint”^17^. Unlike ChIP-seq, which requires a high-quality antibody and can only be run for one TF at a time, ATAC-seq footprints can detect binding sites without specifically knowing the identity of the bound TF. In recent years, multiple algorithms have been developed to quantify binding strength using footprint patterns in ATAC-seq or DNase-seq data, albeit in relatively small sample sizes^18–21^. Footprinting analysis is valuable for implicating causal variants, as it can overcome limitations of resolution in GWAS/eQTL studies due to LD constraints by precisely locating TF binding, and implicate specific TFs based on binding motifs at the footprint location^22^.

Recent studies have investigated the sequence dependency of TF binding via allele-specific cleavage patterns in ATAC-seq^23^, multiplex protein-DNA binding array^24^, or allele-specific ChIP-seq^25^. Most recently, a study used DNase-seq footprints in LCLs from 57 genotyped individuals to uncover SNPs associated with footprint-inferred TF binding, known as footprint QTLs (fpQTLs)^26^. However, this study, and many other footprinting studies^20,21,27^ were limited to sites which overlapped known TF sequence motifs. Given the limited ability of sequence motifs to computationally predict true binding locations^24,28^, this motif-centric approach is much less powered to detect binding events compared to motif-agnostic approaches.

Here, we applied footprinting analysis to a uniformly generated dataset of human liver ATAC-seq samples from 170 genotyped individuals, the largest sample size to date, to measure TF binding strength genome-wide. GWAS have uncovered hundreds of loci associated with liver-related traits including metabolic associated steatotic liver disease (MASLD, formerly NAFLD)^29–31^, type 2 diabetes (T2D)^32^, hyperlipidemia^13,33^, enzyme levels^34^, and T2D risk factors such as obesity^35–37^, most of which have unknown functional mechanisms^38^. We report 693 fpQTLs associated with TF binding at an FDR of 5%. fpQTLs are enriched in transcription start site (TSS)-proximal regions, ChIP-seq peaks for liver-expressed TFs, lipid-associated loci, and molecular QTLs for expression (eQTLs) and chromatin accessibility (caQTLs) mapped in human liver. Notably, the measured effect of an fpQTL on TF binding was highly concordant with its effect on an underlying sequence motif. Finally, we demonstrate that fpQTL discovery can fine-map GWAS loci by pinpointing the causal variant and implicate a specific TF whose binding motif is disrupted. Our map of genotype-dependent TF binding sites offers the opportunity to (1) interpret functional non-coding variants by proposing TF binding as a biological mechanism for association, and (2) aid the identification of the active variant(s) at GWAS loci where the causal variant was previously unknown due to LD-related constraints.

## Materials and Methods

### Study population

The study utilized data collected from 189 specimens obtained from liver transplant recipients from their respective donor cohorts at the University of Pennsylvania, collected in 2012-2017 and 2018-2020 enrolled under the BioTIP study (Biorepository of the Transplant Institute at the University of Pennsylvania). Participants were enrolled in the prospective biorepository and clinical databases, collecting biological samples and clinical data at the time of transplantation, and at predetermined intervals after transplantation. The study was approved by the University of Pennsylvania’s Institutional Review Board (2018-2020: FWA00004028, protocol #814870). All research was conducted in accordance with both the Declarations of Helsinki and Istanbul. The participants signed informed consent forms before transplantation and at the time of organ donation. Specimens collected from this protocol used in this study were deidentified and subsequently anonymized.

### ATAC-seq Library Generation

Human liver wedge biopsies were supplied by the Penn Transplant Institute. Samples were derived from human livers deemed fit for transplantation, and were collected at the time of the surgery. Samples were flash frozen and stored at −80 C. Chromatin accessibility profiles were generated using a modified Assay for Transposase Accessible Chromatin with high-throughput sequencing (ATAC-seq) called Omni-ATAC^16,39^. Briefly, approximately 20 mg of tissue was dounce homogenized in a homogenization buffer. Tissue homogenate was layered over Iodixanol density gradient and spun. Nuclei were extracted post-centrifugation and quantified using a hemocytometer. Approximately 50,000 nuclei were rinsed and added to the Omni-ATAC reaction mix. Transposition reactions were incubated at 37 C for thirty minutes. Reactions were cleaned with spin columns and eluted. Polymerase chain reaction (PCR) was initially performed for five cycles. At this point, a qPCR reaction was performed to determine the additional number of PCR cycles to use. The additional number of PCR cycles was determined by calculating the qPCR cycle at which the fluorescence intensity was equal to one-third the maximum fluorescent intensity of the reaction. Libraries were purified and profiles were measured using Bioanalyzer High-Sensitivity DNA Analysis Kit (Agilent). Libraries that passed visual quality control and concentration checks were frozen at −20 C.

### ATAC-seq Library Sequencing

Libraries were pooled in two separate groups, 93 samples and 96 samples, and sequenced at Vanderbilt University Medical Center (VUMC VANTAGE (Vanderbilt Technologies for Advanced Genomics)) on the Illumina NovaSeq 6000 with PE150 sequencing. Libraries were pooled and sequenced such that each sample was covered by approximately fifty million sequencing reads.

### ATAC-seq Data Processing

ATAC-seq data were processed following the ENCODE processing pipeline, with slight modification. Briefly, FASTQ files were processed with fastp (v.0.12.5) with parameters “-y -c -g”. FastP processed FASTQ files were aligned to GRCh38 using bwa mem (v. 0.7.17-r1188) and piped into samtools (v.1.9) view with parameters “-S -b -f 2 - > outFile.bam” to generate bam files. Duplicate reads were marked and removed using Picard Tools (v.1.141) MarkDuplicates with parameters “ASSUME_SORTED=true, REMOVE_DUPLICATES=true”. Autosomal reads only were retained using samtools view with parameters “input.bam -b {1..22} > ${i}.auto.bam”. Open chromatin peaks were called on all 189 samples using Genrich (v0.6.1) (https://github.com/jsh58/Genrich) with parameters -j, -m 10 and -g 50.

### Genotyping and Imputation from Low Coverage Whole Genome Sequencing

Sample genotype was obtained using low-pass whole genome sequencing from Gencove. Genotypes were filtered to retain only polymorphic sites within our sample population. Polymorphic genotypes were filtered on minor allele frequency (MAF) > 0.05 and genotype posterior probability (GP) > 0.8. Genotypes were phased using Eagle (v2.4.1)^40^.

Of the 189 ATAC-seq samples, 14 were removed for poor genotyping quality, and 5 were removed for having a low read count (< 30 million reads), leaving 170 samples used in fpQTL discovery (see **Supplementary Table 1, Figure S1A** for sample ancestry and covariate information).

### Calculation of footprint scores

Footprint scores were calculated using PRINT^41^ (https://github.com/HYsxe/PRINT, commit 2023-05-14). ATAC-seq bam files were processed into fragment files as described on the PRINT Github (**Web Resources**). Read pairs were removed during this step if (i) both reads mapped to a different chromosome, or (ii) read 1 mapped to the - strand, but did not cover the entire fragment (insertions at these reads did not show the expected Tn5 sequence bias).

Every variant that was found within an open chromatin peak and had MAF > 0.05 within our samples was expanded into a region with 100 bp on either side of the variant (by default, PRINT uses a “context radius” of 100 bp, meaning the outer 100 bp of a region are needed to calculate the background insertion distribution). The Tn5 sequence bias in these regions was calculated by PRINT using the model trained by Hu et al^41^.

For every ATAC-seq sample, getTFBS() was run on the variant regions, following the vignette provided on the PRINT Github (**Web Resources**). The only non-default parameter was tileSize = 1, to measure only the FP score at the variant. The vector of TF binding scores (FP scores) across all variants was extracted for each sample, and these vectors were combined into the footprint score matrix (n = 170, # variants = 3,258,578).

### fpQTL discovery

The distribution of FP scores in each sample was quantile-transformed using the average empirical distribution observed across all samples, following the lead of GTEx *cis*-eQTL mapping^42^. However, the FP scores for each variant were *not* transformed to the quantiles of the standard normal distribution, in order to preserve the signal of extreme FP scores (i.e. FP score ≈ 1)

For every variant considered, the following regression was run in R:

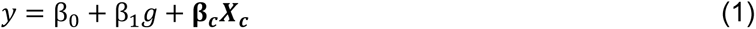

Where *y* is the vector of FP scores across all samples, *g* is the vector of genotypes across all samples (represented by an additive model as the number of non-reference alleles), and ***X_c_*** is a matrix of covariates across samples which includes sex, sequencing batch, and the first three principal components inferred from genotypes. The estimate of *β*_1_ was taken as the estimated fpQTL effect size of the variant, and the fpQTL significance was calculated as the *P*-value of the t-test (two-sided) under the null hypothesis that *β*_1_ = 0. To test the effect of covariates on regression results, we also performed this regression analysis excluding covariates and observed that SNP *P*-values did not change drastically (**Figure S2E**). Multiple test correction was performed using a false discovery rate (FDR) q-value method^43,44^. Variants with a calculated FDR q-value < 0.05 were labeled fpQTLs.

### Allele-specific footprinting

For a given fpQTL, the allelic origin of aligned fragments within heterozygous samples could be determined if at least one of the paired reads overlapped the SNP. Such fragments were separated by allele, and the resulting insertion sites were combined across samples (due to low coverage of allelic fragments) to create “allele-specific insertion patterns”. These insertion patterns were then fed into PRINT separately to calculate allele-specific footprint scores for that fpQTL.

### Gene locations

The locations of transcription start sites (TSSs) were based on NCBI RefSeq’s curated list of genes^45^. The table ncbiRefSeqCurated was downloaded from the UCSC Genome Browser on Jan 22, 2024.

### Liver eQTLs

Association data for expression QTLs from GTEx Analysis V8 were downloaded from the GTEx Portal^8^ (GTEx_Analysis_v8_eQTL.tar)

### Liver caQTLs

caQTLs were called using the same 189 liver samples, as described in a companion manuscript by B.M.W. (in preparation).

### fpQTL overlap with GWAS loci

GWAS summary statistics and lead variants for BMI^36^, T2D^32^, MASLD^31^, enzymes (alanine transaminase/ALT, alkaline phosphatase/ALP, gamma-glutamyl transferase/GGT)^34^, and lipids^33^ were downloaded from their respective publications (**Supplementary Table 5**). LD proxies for lead variants were found using the online tool SNiPA^46^ with Variant Set = 1000 Genomes Phase 3v5, Population = European, and LD (r^2^) threshold = 0.8.

For each liver-related trait, we constructed a 2×2 table, where one dimension represented fpQTLs, and the other dimension represented SNPs which were in LD (r^2^ > 0.8) with a GWAS sentinel variant for that trait. 52805 SNPs, including 20 fpQTLs, did not have LD info in the 1000 Genomes Phase 3v5 SNP set used by SNiPA, and so were not included in the tables. *P*-values and odds ratios were computed using Fisher’s exact test on this table.

### fpQTL enrichment for disease heritability

We performed stratified LD score regression using LDSC (http://www.github.com/bulik/ldsc) v.1.0.1 with the --h2 flag to estimate SNP-based heritability of liver-related traits. We created an annotation consisting of only significant fpQTL SNPs, which was then used to compute annotation-specific LD scores and enrichment for each liver trait.

The baseline model LD scores, plink filesets, allele frequencies and variants weights files for the European 1000 genomes project phase 3 in hg38 were downloaded from the Alkes group (**Web Resources**).

### ChIP-seq peaks

ChIP-seq peaks from ENCODE 3^47^ were downloaded on the UCSC Genome browser (tables encRegTfbsClustered and encRegTfbsClusteredSources) on April 25, 2022. Peaks were considered “liver TF peaks” if at least one of the listed sources was HepG2, liver, or hepatocyte (see **Supplementary Table 3** for list of TFs).

### fpQTL motif matching

Position weight matrices (PWMs) for transcription factor motifs were downloaded from the JASPAR 2024 CORE non-redundant vertebrate database^48^. See **Supplementary Table 4** for list of motifs and corresponding ChIP-seq TFs.

Motif scores were calculated by matching JASPAR motifs to a window around each SNP (for both the reference and alternate allele) using the motifmatchr^49^ (v1.20.0) package, a wrapper for the MOODs^50^ library. SNPs were removed from this analysis if the base in the hg38 sequence did not match the reference or alternate allele from genotyping (n=317 SNPs filtered, remaining # SNPs = 3,258,261). Windows were sized based on motif length, to guarantee that a matched motif would overlap the SNP. We defined an fpQTL-motif overlap to occur when an fpQTL overlapped both a motif and a liver ChIP-seq peak from the corresponding TF.

### Coordinate intersections

All coordinate intersections were calculated in R (v4.4.0) using GenomicRanges (v1.56.0). Coordinates were lifted between hg38 and hg19 as necessary using rtracklayer (v1.64.0).

### fpQTL enrichment with allele-specific ChIP-seq peaks

rsIDs for all fpQTLs were fed into the online tool ANANASTRA^51^, which calculates enrichment for SNPs within the ADASTRA^25^ database for allele-specific ChIP-seq peaks, using the Local (1 Mb) background option.

### Calculation of Tn5 insertion density

To assess the general enrichment in chromatin accessibility near the ends of chromosomes, we divided the genome into bins with a width of 200 bp (the width that PRINT uses to calculate FP score). For each sample, we counted the number of Tn5 insertions in each bin using the fragment file. The first and last 10,000 bins on each chromosome (i.e. within 2 Mb of a chromosome end) were labeled “near-telomeric” bins, and the other bins were labeled “central”. We then averaged the number of insertions in non-empty near-telomeric and central bins to calculate the mean insertion density for these two regions.

### Inversion haplotyping

We used the scoreInvHap^52^ (v1.20.0) R package (https://rdrr.io/bioc/scoreInvHap/) to call inversion haplotypes using the genotypes of nearby SNPs. Data for the inversions called are included in the package.

### Correcting for Tn5 insertion bias when visualizing Tn5 insertions

For every variant in every sample, PRINT considers the Tn5 insertions at the *x*-th position within a 200 bp window around the variant. If Ox represents the observed cut sites at the *x*-th position in the sample, then we calculated the corrected cut sites C_x_ in that sample as:

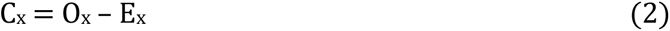

where E_x_ is the expected number of cutsites at position x calculated by:

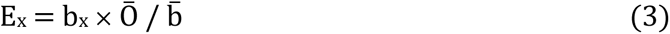

where bx is the Tn5 bias at the *x-*th position reported by PRINT, and 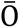 and 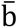 are the means of O_x_ and b_x_ calculated across all *x* positions within the 200 bp window.

## Results

### Discovery of liver fpQTLs in liver open chromatin

Using a human liver ATAC-seq dataset uniformly generated from 170 genotyped donors (**Figure S1A**), we first scored all common SNPs residing within open chromatin regions for their potential of TF footprints by calculating a footprint score (FP score) for each sample at every SNP position (**Materials and Methods, Figure 1A**). For each SNP, we then performed linear regression analysis to estimate the effect of change in genotype on FP score as the outcome, including covariates (**Materials and Methods**, **Figure 1B**). 693 SNPs exceeded an FDR q-value < 0.05 multiple-testing threshold and were labeled as fpQTLs (**Figure 2A, Supplementary Table 2**).

**Figure 1.**
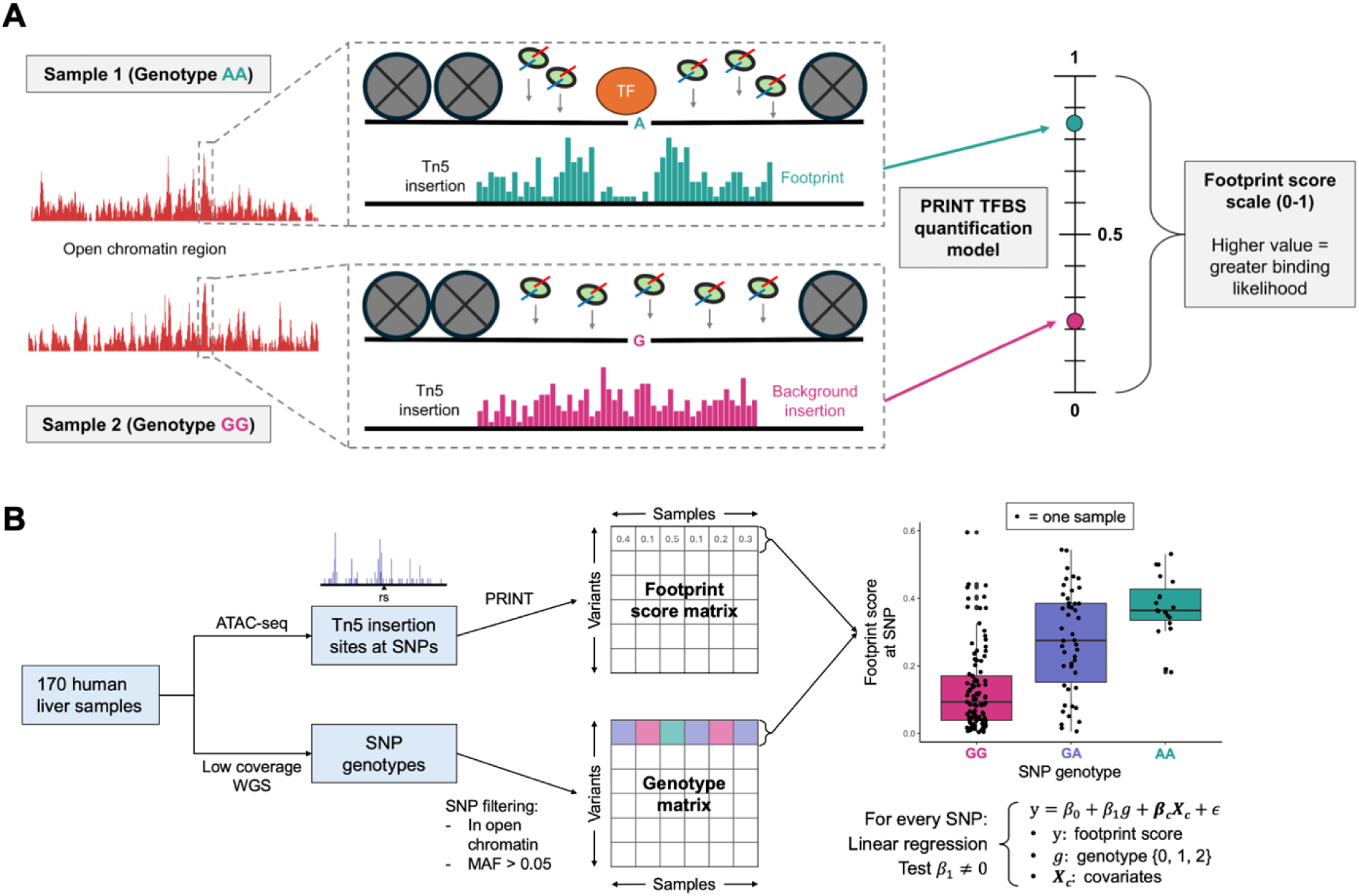
ATAC-seq footprinting analysis can detect genotype-dependent binding events. **(A)** Calculation of FP score. TF binding is detectable in ATAC-seq experiments because bound TFs block the insertion of Tn5, leaving a site of relatively depleted cutsites within a larger ATAC-seq peak, known as a footprint. The PRINT software calculates the footprint (FP) score of a local insertion pattern using a supervised regression model trained on the insertion patterns of known binding sites. The resulting FP score can be interpreted as the relative likelihood of a binding event, which can depend on the genotype of a local SNP. **(B)** fpQTL discovery. Liver samples were taken from 170 donors, and analyzed by ATAC-seq and whole-genome sequencing (WGS). PRINT was used to calculate a footprint score at every SNP location in every sample, and for every SNP an FP score was regressed onto SNP genotype across samples to calculate a *P*-value for the strength of association.

**Figure 2.**
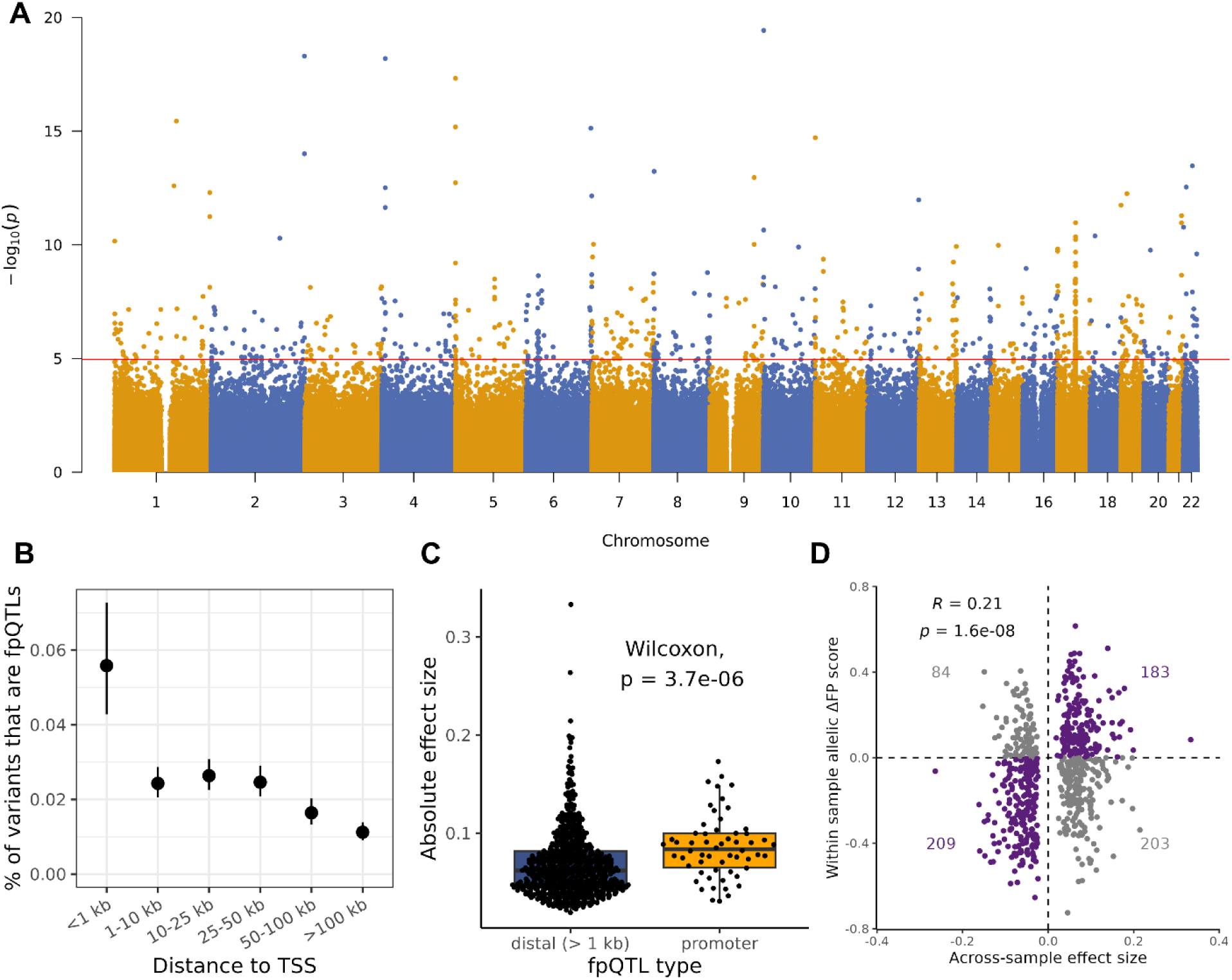
fpQTLs are enriched near transcription start sites (TSSs) **(A)** Manhattan plot. For each SNP, the FP score was regressed onto genotype to calculate a coefficient of association (β_1_). *P*-values were calculated by testing the null hypothesis that β_1_ = 0. The vertical line represents an FDR-adjusted *P*-value of 0.05. **(B)** fpQTL proportions based on TSS-proximity. SNPs were binned based on distance to the nearest TSS, and the proportion of SNPs within the bin labeled as fpQTLs was calculated. 95% binomial confidence intervals are shown. **(C)** Promoter fpQTLs have higher effect sizes (|β_1_|) than TSS-distal fpQTLs. fpQTLs were considered within a promoter if the distance to the nearest TSS was < 1 kb. **(D)** For all fpQTLs, the regression β_1_ (x-axis) is plotted against ΔFP score = alt allelic FP score – ref allelic FP score (y-axis), where the allelic FP scores were calculated by considering insertions in heterozygous samples separately based on their allele. Purple fpQTLs are concordant between their across sample and within-sample effect. The number of fpQTLs is labeled in each quadrant (Fisher’s exact test OR = 2.2, *P*-value = 8.1×10^-7^).

When comparing fpQTL positions to the locations of gene transcription start sites (TSSs), we observed that TSS-proximal SNPs were more likely than distal SNPs to be detected as fpQTLs, and TSS-proximal fpQTLs had higher effect sizes than distal fpQTLs (**Figures 2B and 2C**). It has been shown previously that proximal loci are more likely to be in highly accessible chromatin and to have regulatory significance^53^, suggesting greater power to detect fpQTLs in TSS-proximal regions. As such, we reasoned that our observed enrichment was likely due to greater statistical power to detect, rather than proximal SNPs having a stronger effect on TF binding.

To account for systematic biases in ATAC-seq insertion across samples, we next considered the differences in insertion patterns on each allele within heterozygous samples. We predicted that for true fpQTLs, the allele associated with increased binding within heterozygous samples would also be associated with FP score across all samples. For each fpQTL, we calculated allelic footprint scores using insertions from the reference and alternate alleles separately within heterozygous samples (**Materials and Methods**). We observed that the difference between alternate and reference footprint scores was significantly correlated with fpQTL effect size in the expected direction (**Figure 2D**).

### fpQTL effects are concordant with TF binding motifs

For TFs with high-quality antibody availability, binding sites can be mapped with very high confidence using chromatin immunoprecipitation sequencing (ChIP-seq) data^54^. To assess the accuracy of our footprint binding detection, we compared the locations of our fpQTLs to known binding locations from liver ChIP-seq data in ENCODE^47^ (**Materials and Methods**). After correcting for multiple testing, we observed that fpQTLs were over-represented in liver ChIP-seq peaks for 49 of the 121 TFs with available data. In particular, we observed enrichment for several TFs known to be associated with liver metabolism and disease, including HNF4A^55^ (OR = 4.1, *P* = 6.6×10^-16^), FOXA1 (OR = 2.6, *P* = 3.7×10^-5^), FOXA2^56^ (OR = 2.6, *P* = 1.5×10^-5^), and JUND^57^ (OR = 3.5, *P* = 1.1×10^-10^) (**Supplementary Table 3**, **Figures 3A and S3A**).

**Figure 3.**
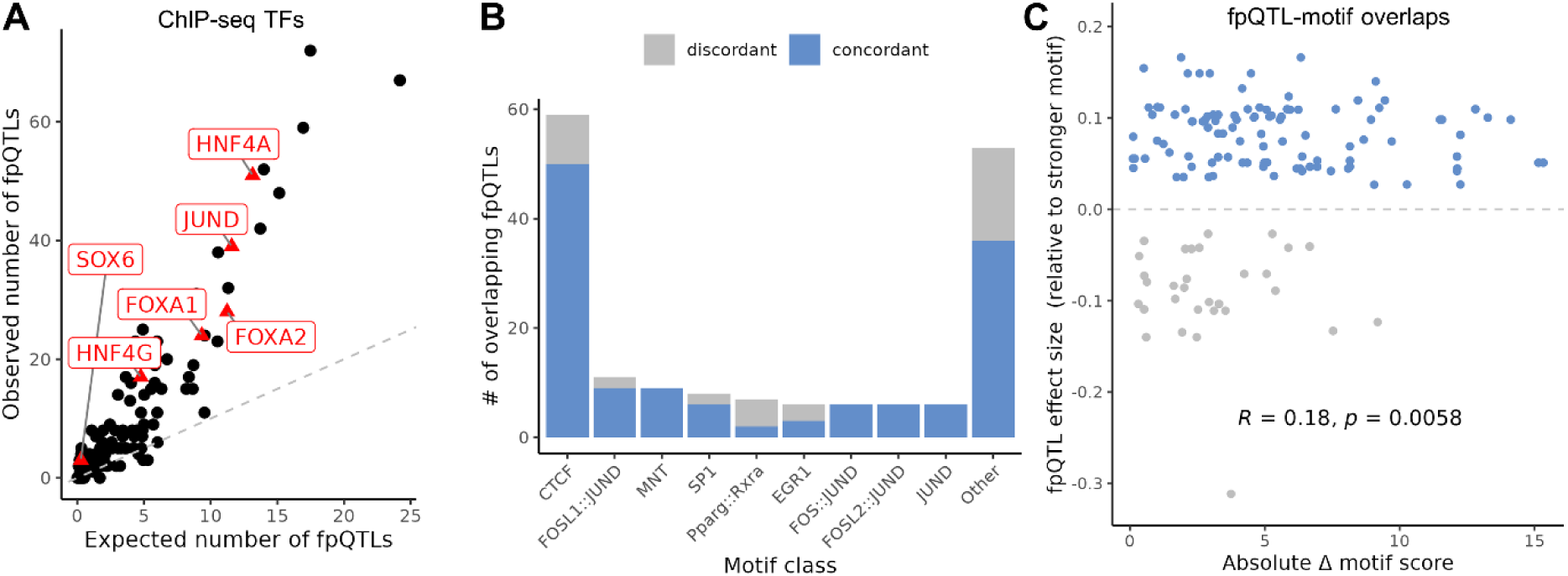
fpQTLs are enriched in ChIP peaks and concordant with underlying sequence motifs. **(A)** The expected and observed number of fpQTLs within ChIP peaks for every TF with ChIP data. Liver-related TFs are labeled in red (see **Supplementary Table 3**). Expected number of fpQTLs was calculated as [#SNPs in ChIP peaks × proportion of SNPs that are fpQTLs]. **(B)** Number of concordant and discordant fpQTLs which overlap given motifs, grouped by TF. Three redundant CTCF motifs were excluded. Motifs from JASPAR, matched with *P*=5×10^-4^. **(C)** Comparison of fpQTL effect size with the change in motif score, for all fpQTL-motif overlaps. The y-axis represents the regression beta, with positive values indicating an increase in binding for the allele with the stronger motif. Spearman coefficient and *P*-value shown.

We next examined how fpQTLs altered the strength of binding motifs underlying these ChIP-seq peaks. Specifically, we assessed whether fpQTLs were “concordant” with the motifs they overlapped; that is, if the allele with the stronger motif match was associated with a higher FP score^25^. Among the fpQTL-motif overlaps that we identified at ChIP-seq peaks, a large proportion (181/227, 80%) were concordant (**Supplementary Table 4**, **Figure 3B**; binomial *P*-value < 2.2×10^-16^). Additionally, increasing the motif matching significance (*P*-value) threshold by a factor of 10 increased this concordance proportion to 91% (**Figure S4**). Furthermore, the allelic change in motif score was significantly correlated with the FP score-inferred change in TF binding (**Figure 3C**, Spearman’s rho = 0.18, *P*-value = 5.8×10^-3^), suggesting that variants with a larger impact on a given motif are more likely to be concordant.

To test our hypothesis that fpQTLs represent allele-specific binding, we compared our fpQTLs to SNPs which showed allele-specific ChIP-seq peaks in the ADASTRA^25^ database. We found that our set of fpQTLs were significantly enriched for SNPs with allele-specific binding in ChIP-seq for HepG2 cells and liver (**Figure S3C**), suggesting that fpQTLs reflect true allele-specific binding effects. Taken together, these results suggest that fpQTLs influence TF binding strength by disrupting the binding sequence motif for the given TF.

### fpQTLs are significantly enriched in other liver QTLs and lipid GWAS signals

To assess the role of fpQTLs in regulating gene expression, we investigated the overlap of fpQTLs with genetic variation associated with expression in liver (i.e., liver eQTLs) from GTEx^8^, and caQTLs discovered using the same set of liver samples studied here. We observed a highly significant enrichment of fpQTLs for liver eQTLs (odds ratio = 4.01, P = 1.1×10^-20^), and an even higher enrichment for caQTLs (odds ratio = 29.3, P < 2.2×10^-308^; **Supplementary Table 5**). For the set of n=76 SNPs that were both fpQTLs and eQTLs, the allele associated with increased TF-binding was also associated with increased gene expression in 55 (72%) SNPs (**Figure S5A**). This suggests that most of the regulatory elements harboring fpQTLs act as enhancers of gene expression, where TF-binding promotes transcription, rather than as silencers. An even stronger directional correspondence was observed for chromatin accessibility, where 344 (93%) fpQTL/caQTLs act in the same direction, compared to only 27 in the opposite direction (**Figure S5B**).

We next queried if fpQTLs can account for a proportion of the characterized genetic component of disease. We examined the overlap of fpQTLs with lead SNPs and their LD proxies identified by liver-related GWAS (**Materials and Methods**). fpQTLs were significantly enriched for GWAS SNPs for four out of five lipid traits; however, we did not observe any fpQTLs overlapping with GWAS SNPs for BMI, T2D, or MASLD (**Supplementary Table 5**, **Figure 4A**). However, fpQTLs were not depleted for any trait, suggesting that the lack of overlap is in part driven by the small number of fpQTLs (i.e., low statistical power).

**Figure 4.**
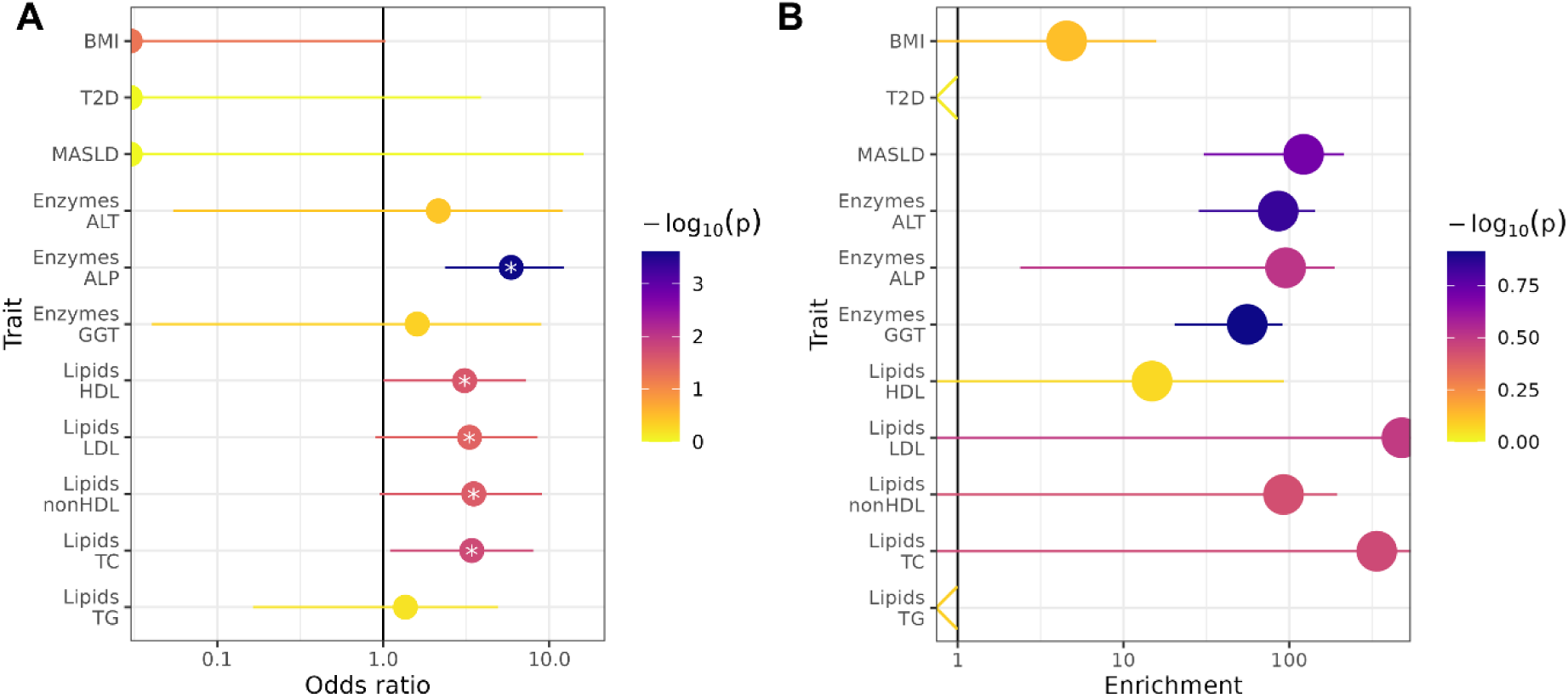
fpQTLs are enriched for lipid-associated SNPs. **(A)** Enrichment of fpQTLs in GWAS/QTL SNPs for different traits, using odds ratios (OR). GWAS SNPs investigated were defined as all SNPs which are either (1) a lead SNP reported in literature, or (2) a proxy of a lead SNP with r^2^ > 0.8. The top three traits have no such GWAS SNPs as an fpQTL (OR = 0). *P*-values come from Fisher’s exact test, 95% confidence intervals are shown. Traits which are nominally significant (P < 0.05) are annotated with **\***. **(B)** Enrichment of GWAS heritability in fpQTLs for several traits, calculated by stratified LD score regression. *P*-values are calculated by ldsc using permutations. Error bars show ± standard error of enrichment. ldsc can sometimes return negative enrichment values, which are indicated for T2D and TG.

Given that odds ratio enrichment tests do not take into account the LD structure of SNP associations, we also evaluated enrichment for disease heritability at fpQTLs using stratified LD score regression^58^. Despite observing very high estimated heritability enrichment for MASLD and lipid traits (**Figure 4B**), none of these tests were statistically significant given the fpQTL annotation was relatively small compared to the entire genome. However, the high heritability enrichment in traits without a high overlap with lead SNPs or proxies suggests that fpQTLs overlap several sub-significant signals which the GWAS were not powered to detect. Overall, the enrichment of fpQTLs with GWAS signals was much lower compared to enrichment with other forms of QTL.

### Greater chromatin accessibility increases power to detect fpQTLs

We also observed that the likelihood of a SNP being labelled as an fpQTL increased with the degree of openness of its chromatin peak. Indeed, fpQTLs are located within peaks with a higher average openness across samples (measured in ATAC-seq fragment counts per million, or CPM) compared to non-fpQTL SNPs (**Figure S6A**), and the measured effect size and significance of fpQTLs is significantly correlated with the average number of Tn5 insertions near the fpQTL (**Figures S6B and S6C**). We hypothesize that this effect is due to both (i) enrichment of functional significance in highly accessible regions^58–60^, and (ii) greater power to detect footprint activity when PRINT can consider more Tn5 insertions to calculate FP score. For example, if a TF-bound SNP is located in a relatively inaccessible peak, then the lack of local insertions will prevent PRINT from confidently assigning a high FP score, despite the presence of a TF.

We next tested whether chromatin accessibility increases fpQTL power or if this effect is driven solely by increased functional activity within highly accessible peaks. We elected to measure the relationship between the average number of insertions near a SNP and its average FP score across samples. We observed that high insertion counts corresponded to higher FP scores even after removing SNPs with known regulatory function (**Figures S6D-F**). This observation suggests that PRINT skews towards assigning higher FP scores to SNPs near more insertions, regardless of functional activity.

The increase in fpQTL power due to chromatin accessibility explains our observation that fpQTLs are enriched nearer the ends of chromosomes, closer to the telomeres. Of SNPs within 2 Mb of the chromosome ends, 130 (0.08%) are fpQTLs, compared to 563 (0.02%) for the more central SNPs (odds ratio = 4.3, Fisher’s *P*-value < 2.2×10^-16^). These “telomeric-neighboring” regions were in fact not enriched for the ChIP-seq peaks of any TF, which therefore did not explain the high concentration of fpQTLs at these locations. Instead, we hypothesize that telomeric-neighboring regions are enriched for fpQTLs because they are more accessible on average than other regions, therefore increasing fpQTL detection power. Indeed, telomeric-neighboring regions (within 2 Mb of chromosome ends) had significantly higher insertion density compared to central regions in all of our liver ATAC-seq samples (**Figure S7A**). fpQTLs in these telomeric-neighboring regions resided in peaks with a significantly higher mean CPM and were flanked by a higher number of Tn5 insertions (**Figures S7B and S7C**).

In addition to an enrichment of fpQTLs near the ends of chromosomes, the fpQTL Manhattan plot showed a highly significant signal on chromosome 17, which we mapped to a known common inversion at the 17q21.31 locus^61^. Calling inversion haplotypes on our samples and including them in the regression revealed that the significant SNPs on the ends of the inversion were tagging the inversion itself, potentially altering TF binding at its boundary regions. Additionally, this correction revealed four SNPs within the inversion associated with TF binding (**Figure S8**). Indeed, this inversion has been implicated in HDL lipid levels and other obesity related traits^62^, along with brain morphology and neuroticism^63,64^.

### fpQTLs can be used to fine-map GWAS loci

Unlike GWAS and other forms of QTLs, the trait tested against each SNP in fpQTL discovery (footprint score) is different at every SNP, meaning test statistics between nearby SNPs are not correlated due to LD. As a result, fpQTLs provide single-SNP resolution to fine-map GWAS loci or QTLs by pinpointing a putatively causal SNP among a credible set of GWAS/QTL variants. To explore the fine-mapping ability of fpQTLs, we examined GWAS loci for liver traits harboring at least one significant fpQTL.

First, we examined the lipid-associated *SORT1* locus, which has been extensively studied through experimental validation to determine that the association is driven specifically by rs12740374, creating a TF binding site^65,66^. Our fpQTL results identified the same SNP as the causal variant, given rs12740374 was the most significant at the locus (**Figures 5A and 5B**). Assessing the bias-corrected insertions around rs12740374 (see **Materials and Methods** for insertion corrections), samples with the alternate allele showed a strong depletion of insertions directly adjacent to the variant, suggesting a genotype-dependent binding event consistent with previous results^65^ (**Figure 5C**). This rediscovery of a known causal SNP supports the hypothesis that fpQTL discovery can be a powerful tool in fine-mapping.

**Figure 5.**
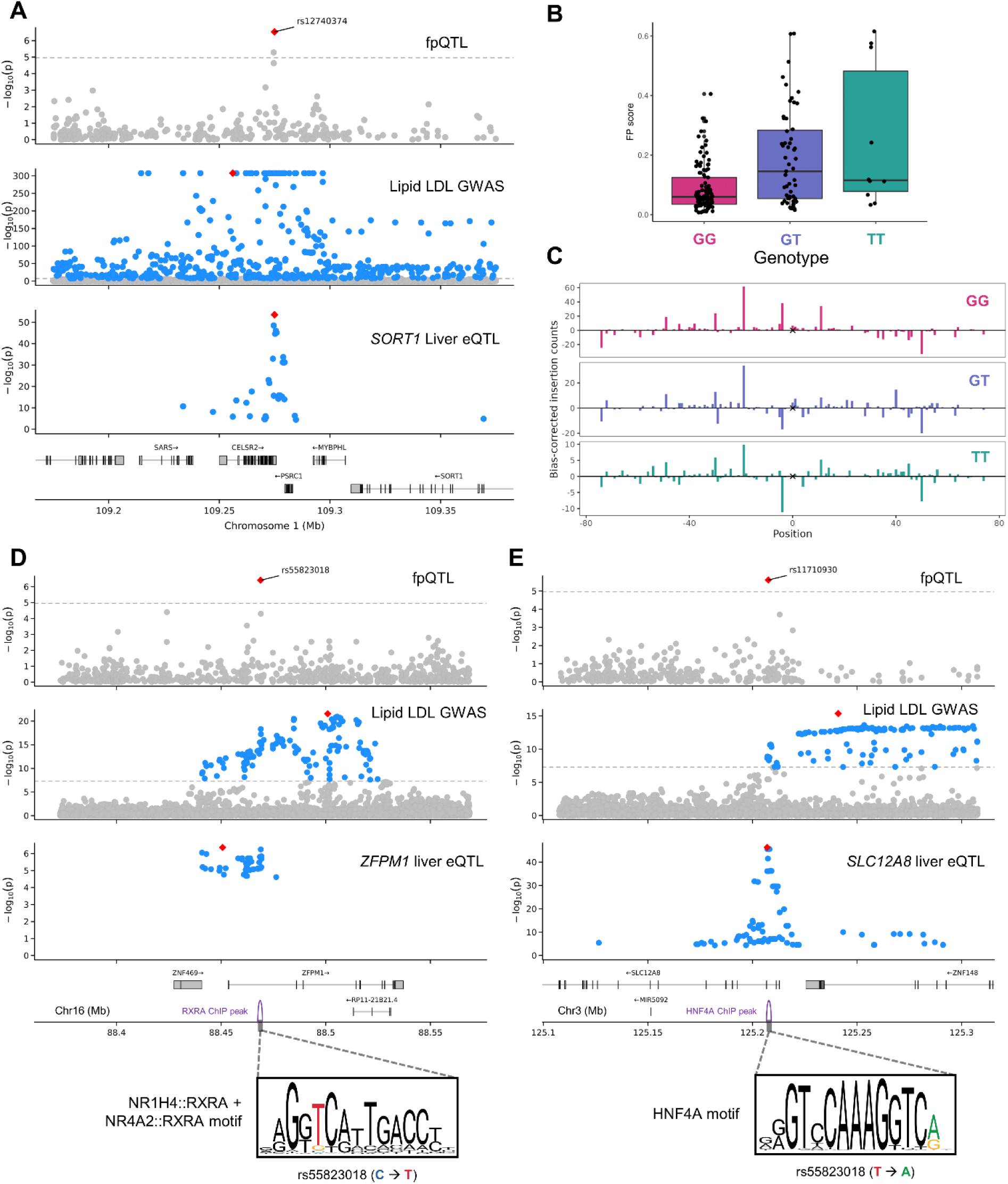
fpQTLs can fine-map GWAS loci. Significance plots show *P*-values for fpQTLs (top), LDL GWAS (middle), and eQTLs (bottom). **(A)** *SORT1* locus significance plot. **(B)** FP score at rs12740374 (at *SORT1* locus) across samples based on genotype. **(C)** Bias-corrected Tn5 insertions around rs12740374 (marked with x) based on genotype, aggregated across samples. **(D)** *ZFPM1* locus significance plot, with the effect of rs55823018 on the RXRA binding motif shown below. **(E)** *SLC12A8* locus significance plot, with the effect of rs11710930 on the HNF4A binding motif shown below

We next sought to investigate fpQTLs which could explain less defined GWAS loci. At the lipid (LDL)-associated *ZFPM1* locus, the fpQTL rs55823018 was by far the most significant compared to adjacent SNPs in partial LD (**Figure 5D**). This SNP increased TF binding and resided in a ChIP-seq peak for the Retinoid X receptor alpha (RXRA), and concordantly increased the matching strength of the underlying NR1H4::RXRA sequence motif. A second instance was observed at the lipid-associated *SLC12A8* locus, where the fpQTL rs11710930 overlapped a ChIP-seq peak for hepatocyte nuclear factor 4 alpha (HNF4A) and was concordant for the underlying motif (**Figure 5E**). HNF4A is well-established as an important TF for liver function, and has been previously implicated in liver dysregulation^55,67–69^. Furthermore, *SLC12A8* has been implicated as an effector gene for T2D risk^70^, but the role of this locus in lipid levels remains to be investigated. Taken together with orthogonal ChIP-seq and eQTL results, our fpQTL method adds to the confluence of evidence implicating this specific variant as causal for increased lipid levels.

## Discussion

A significant limitation to GWAS and QTL studies is the unwieldy number of candidate causal variants due to constraints of LD, making it challenging to pinpoint which among them is truly causal. Here, we leveraged statistical inference of TF binding likelihood from experimental data at base-pair resolution to discover fpQTLs, i.e., variants associated with TF binding. We showed that liver fpQTLs were concentrated in ChIP-seq peaks, eQTLs, caQTLs, and lipid-associated loci. Additionally, the vast majority of fpQTLs were concordant with underlying sequence motifs, increasing our confidence that fpQTLs represent SNPs that are very likely to be causal for TF binding differences. We also observed specific examples of GWAS loci where fpQTL discovery implicated both a causal variant and a corresponding disrupted TF binding motif.

The main limitation of this study was the high level of noise in ATAC-seq insertion positions, resulting in high variance in footprint score despite our large read count. Additionally, we used ATAC-seq data from bulk tissue samples rather than single-cell samples, which may mask footprint signals that only occur in a specific cell type. However, given the majority (60%) of liver cells are hepatocytes^71^, we are likely capturing most of the footprint signals from hepatocytes without introducing false signals from mixed cell types. Furthermore, our fpQTL discovery did not consider the effect that each SNP could have on the Tn5 sequence bias, which weakly influences the positions where Tn5 is inserted^16,72^. PRINT corrects for this Tn5 sequence bias, but relies on the reference genome, and so the bias of the alternative allele is not considered. Another limitation of this fpQTL discovery effort was the lack of enrichment in several relevant GWAS traits, limiting their ability to explain disease associations. We propose that this is due to systematic differences between QTL and GWAS discovery. A recent study showed that compared to eQTL signals, GWAS signals are further away from transcription start sites and tend to be near genes under strong selective constraint with more complicated regulatory landscapes^10^. This published model suggests that regulatory variants targeting genes with large trait effects (detected by GWAS) will be less frequent due to natural selection, therefore reducing QTL detection power. Under this model, fpQTL discovery would be similarly hindered at genes with large trait effects, leading to less GWAS enrichment.

Curiously, all of our ATAC-seq samples showed greater Tn5 insertion density in telomeric-neighboring regions compared to other regions. This warrants further investigation to (1) assess the consistency of this phenomenon across cell types and experimental parameters, (2) understand the implication for this bias in peak-calling and multiple-testing correction due to changes in power, and (3) determine the source of this bias, whether technical or biological. However, we deemed these questions outside the scope of our current investigation.

Our results demonstrate that fpQTL characterization enables the capture of genetically regulated TF binding signals in human liver with a resolution not constrained by LD patterns. The approach therefore has the potential to identify regulatory SNPs among trait-associated loci from GWAS or QTL studies, which typically harbor many variants in LD with the causal SNP. fpQTLs also suggest transcription factor binding as the mechanism by which non-coding GWAS variants affect disease risk. Overall, our map of genotype-dependent TF binding sites is a valuable resource for understanding the genetic etiology of complex traits in the context of liver. By implicating specific regulatory elements in these liver-related traits, our fpQTL discovery method should improve research aimed at developing novel therapies by prioritizing variants and TFs for further experimental study. Furthermore, this method can be equally applied to other tissues and cell types, expanding the number of genetic traits that can be addressed.

## Acknowledgments

We thank all members of the Grant and Almasy labs for their feedback on this project. We thank Matt Pahl and Khanh Trang for their assistance running ldsc. We thank Iain Matheson for feedback on this manuscript as well as for sitting on the thesis committee of M.F.D., along with Klaus H. Kaestner and Alexis Battle. We would especially like to acknowledge the mentorship of Christopher D. Brown, who passed soon after this study began, and without whom it would not have been possible. M.F.D. is supported by the National Science Foundation Graduate Research Fellowship Program (NSF GRFP). C.D.B. is funded by R01 HL133218. L.A. is funded by NIAAA U10 AA008401. B.F.V. gratefully acknowledges support from the NIH/NIDDK (UM1 DK126194 and U24 DK138512). S.F.A.G. is funded by UM1 DK126194, R01 HD056465 and the Daniel B. Burke Endowed Chair for Diabetes Research.

## Author contributions

C.D.B. and M.F.D. conceptualized the method. B.M.W. generated and processed the ATAC-seq libraries. M.F.D. performed the data analysis and wrote the manuscript. B.M.W., L.A., and S.F.A.G. reviewed the manuscript and contributed to the statistical methodology. C.D.B is credited posthumously; all other authors read and approved the final manuscript.

## Web Resources

PRINT: https://github.com/HYsxe/PRINT

- Fragment extraction: https://github.com/HYsxe/PRINT/issues/6
- Vignette: https://github.com/HYsxe/PRINT/blob/main/analyses/BMMCTutorial/BMMCVignette.pdf

Genrich (v0.6.1): https://github.com/jsh58/Genrich

GTEx Portal (v8): https://gtexportal.org/home/downloads/adult-gtex/overview JASPAR (2024): https://jaspar.elixir.no

SNiPA (v3.4): https://snipa.org/snipa3/

LDSC (v1.0.1): http://www.github.com/bulik/ldsc

- Data files download: https://console.cloud.google.com/storage/browser/broad-alkesgroup-public-requester-pays/LDSCORE

ANANASTRA (Bill Cipher v5.1.3): https://ananastra.autosome.org/

## Data and code availability

Raw sequencing files for the ATAC-seq samples used in this study have been uploaded to GEO and will be available upon publication. The FP score matrix, genotype matrix, and full fpQTL summary statistics have been uploaded to Zenodo [pending publication, available upon reasonable request]. All intermediate files used in this analysis are available upon request. All code used to process the ATAC-seq samples, run fpQTL discovery, and produce the figures in this manuscript are available on GitHub at https://github.com/maxdudek/fpQTL.

## Supplementary Figures

**Figure S1.**
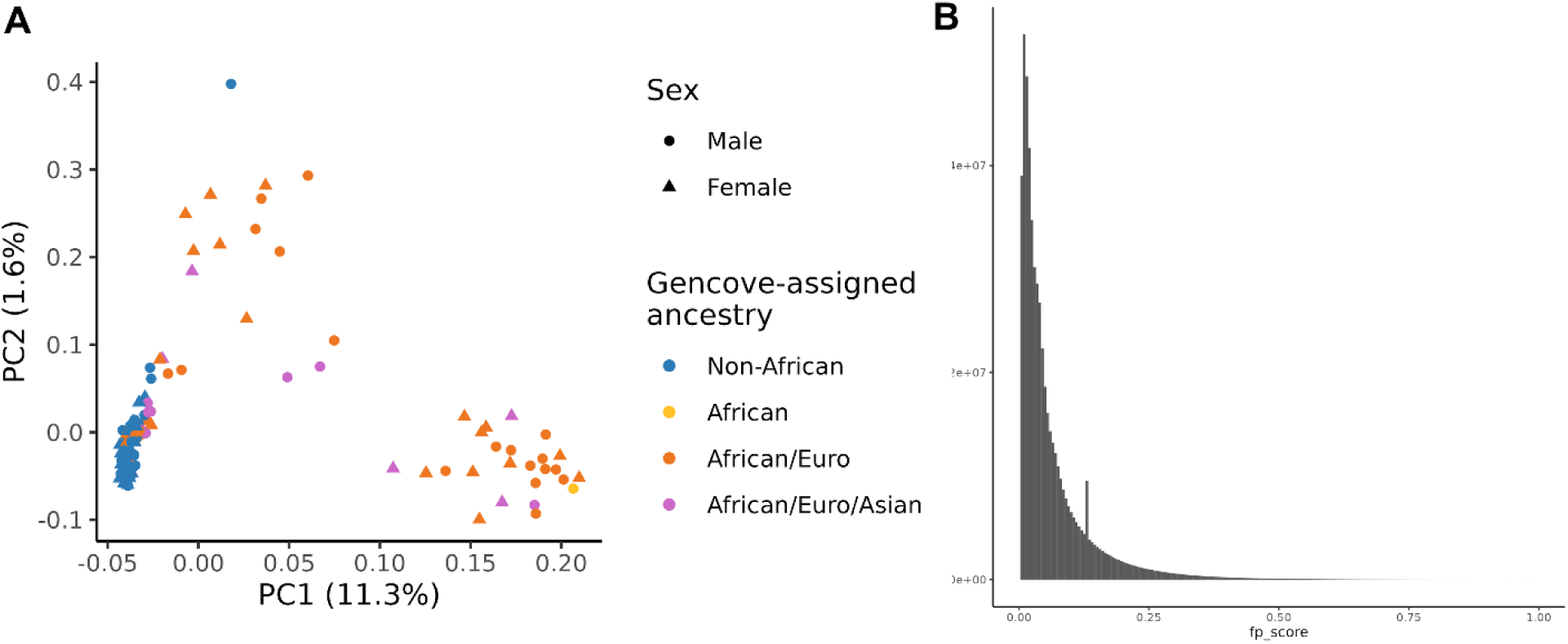
**(A)** 170 ATAC-seq samples plotted along the first two principal components of genotype. The proportion of variance explained by each component is shown on the axes labels. The Gencove genotyping results place each sample into one of four ancestry categories, which are labeled by color. **(B)** Distribution of FP scores across all samples and variants. The spike corresponds to variants in samples with no insertions within a 200 bp window, which are all assigned the same FP score by PRINT.

**Figure S2.**
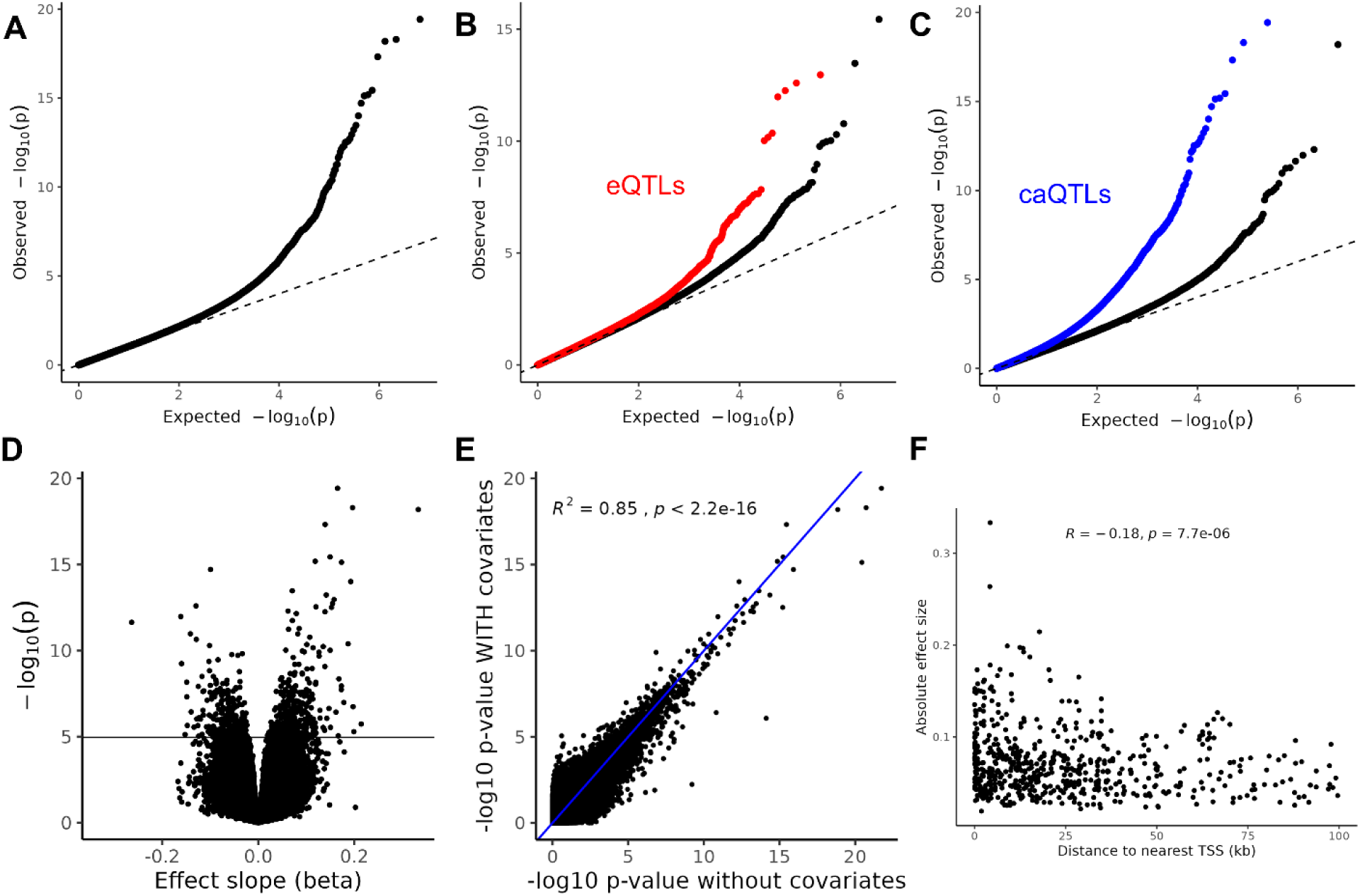
**(A)** QQ plot of fpQTL discovery. **(B)** QQ plot of fpQTL discovery separated based on liver eQTL status. SNPs in red are significantly associated with the expression in liver of at least one gene. **(C)** QQ plot of fpQTL discovery separated based on liver caQTL status. SNPs in blue are significantly associated with the chromatin accessibility of at least one peak, using the same liver samples as fpQTL discovery. **(D)** Volcano plot of fpQTL discovery, with FDR 5% threshold shown. **(E)** Comparison of *P*-values calculated using regressions with and without covariates included. **(F)** Correlation between TSS distance and absolute effect size (|β_1_|) for fpQTLs.

**Figure S3.**
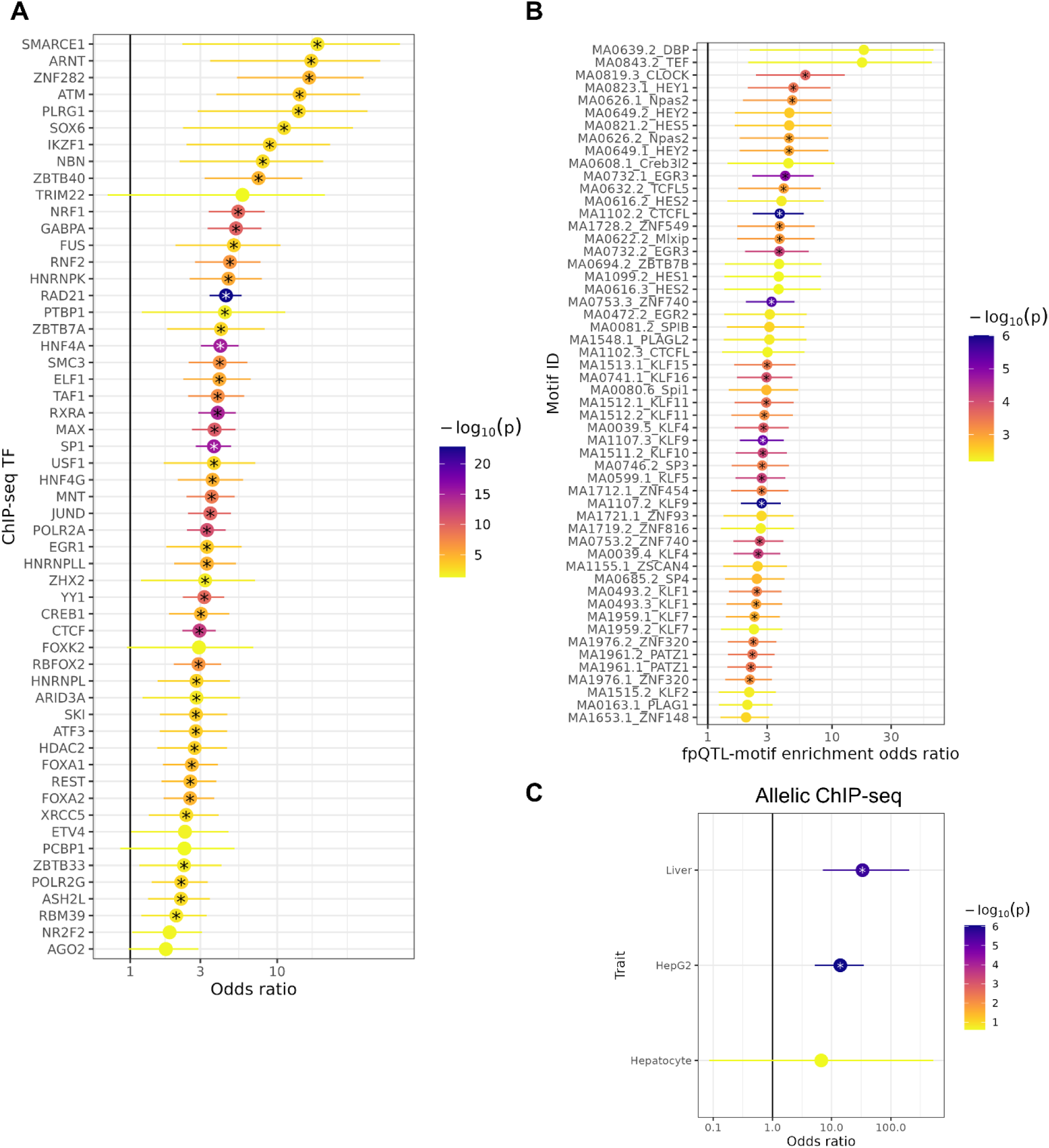
**(A)** Enrichment of fpQTLs in ChiP-seq peaks for different TFs. *P*-values are from Fisher’s exact test on the null hypothesis that the true odds ratio is 1. Tests that passed an FDR-adjusted *P*-value threshold of 0.05 are marked with an asterisk. 95% confidence intervals are shown. See **Supplementary Table 2** for full list of TFs. **(B)** Enrichment of fpQTLs in motif sites, for motifs which do not have corresponding ChIP-seq data. *P*-values are from Fisher’s exact test on the null hypothesis that the true odds ratio is 1. Tests that passed an FDR-adjusted *P*-value threshold of 0.05 are marked with an asterisk. 95% confidence intervals are shown. Motifs matched with *P*=5×10^-5^. **(C)** Enrichment of fpQTLs with allele-specific ChIP-seq peaks from ADASTRA, in 3 liver-related tissue types. *P*-values come from Fisher’s exact test, 95% confidence intervals are shown.

**Figure S4.**
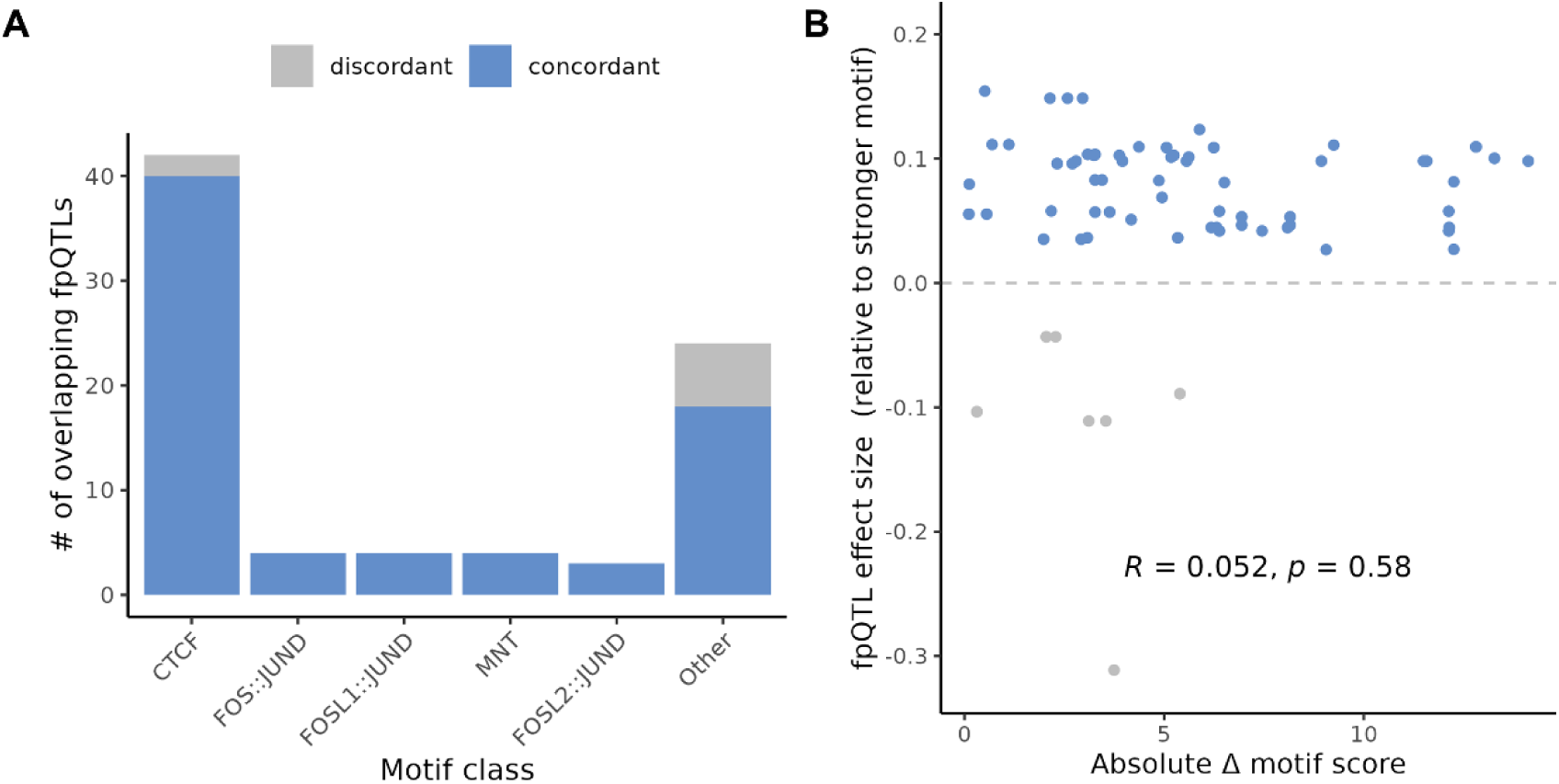
**(A)** Number of concordant and discordant fpQTLs which overlap given motifs. Motifs from JASPAR, matched with *P*=5×10^-5^. **(B)** Comparison of fpQTL effect size with the change in motif score, for all fpQTL-motif overlaps. The y-axis represents the regression beta, with positive values indicating an increase in binding for the allele with the stronger motif. Motifs matched with *P*=5×10^-5^. Spearman coefficient and *P*-value shown.

**Figure S5.**
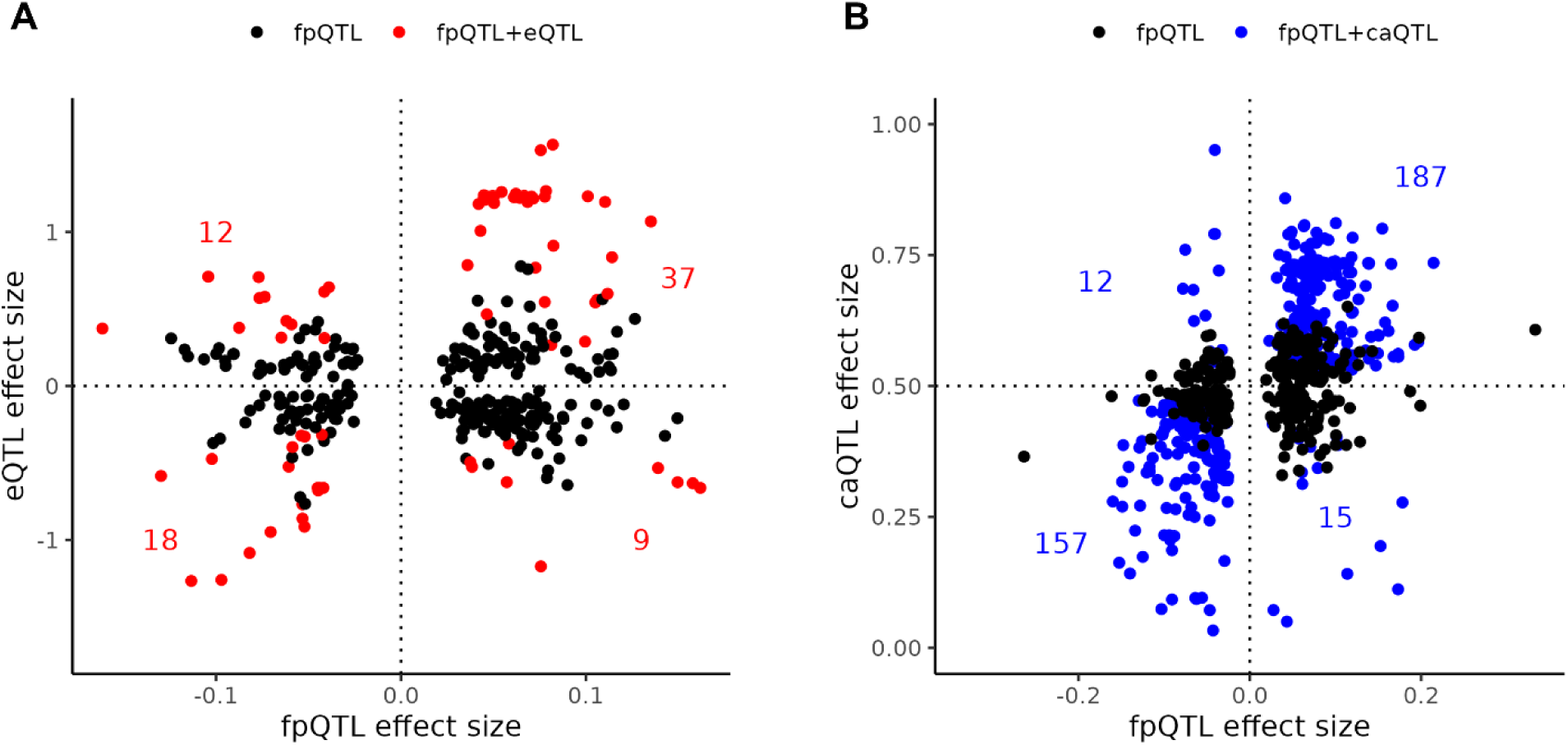
**(A)** Footprint score effect size of fpQTLs compared to eQTL effect size. For every fpQTL, the eQTL effect size for the eGene with the most significant association was used. SNPs that are also significant eQTLs are plotted in red. The number of eQTLs in each quadrant is labeled. **(B)** Footprint score effect size of fpQTLs compared to caQTL effect size (from Rasqual). For every fpQTL, the caQTL effect size for the peak with the most significant association was used. SNPs that are also significant caQTLs are plotted in blue. The number of caQTLs in each quadrant is labeled.

**Figure S6.**
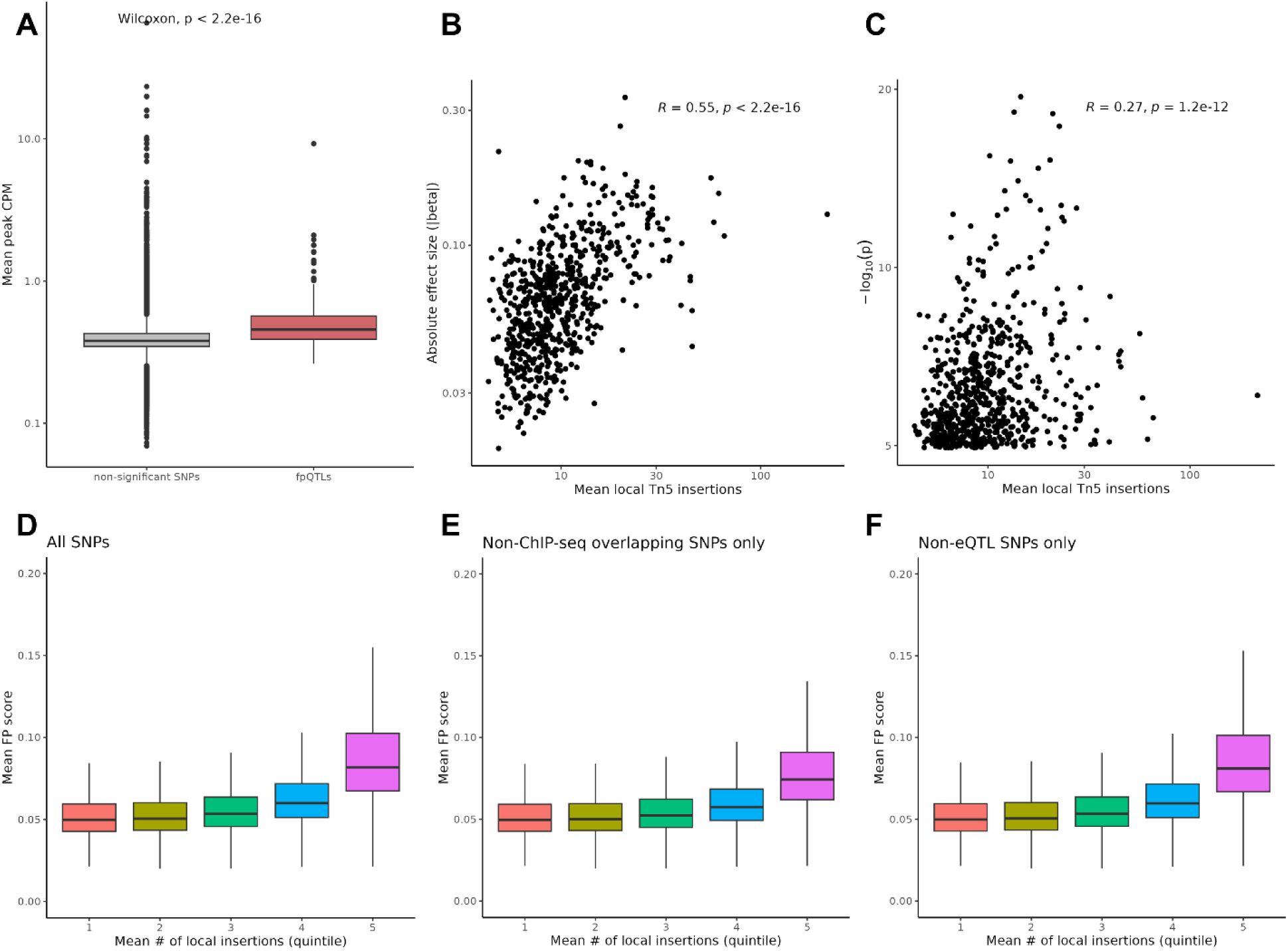
**(A)** Average CPM of a SNPs occupied peak across samples, for both fpQTLs and non-significant SNPs. **(B,C)** The mean number of local Tn5 insertions for fpQTLs (number of insertions within 100 bp, the window used by PRINT to calculate FP score) across samples, compared with (B) the absolute value of the regression beta (effect size), or (C) the -log10 *P*-value from regression. Spearman correlation coefficients and *P*-values from the correlation test are shown. **(D,E,F)** SNPs were placed into quintiles based on the average number of local (within 100 bp) Tn5 insertions across samples, and the distribution of average FP score across samples is shown for each quintile. This was done for (D) all SNPs, (E) excluding SNPs that overlapped a ChIP-seq peak in a liver cell type, and (F) excluding SNPs which were significant liver eQTLs, (**Materials and Methods**).

**Figure S7.**
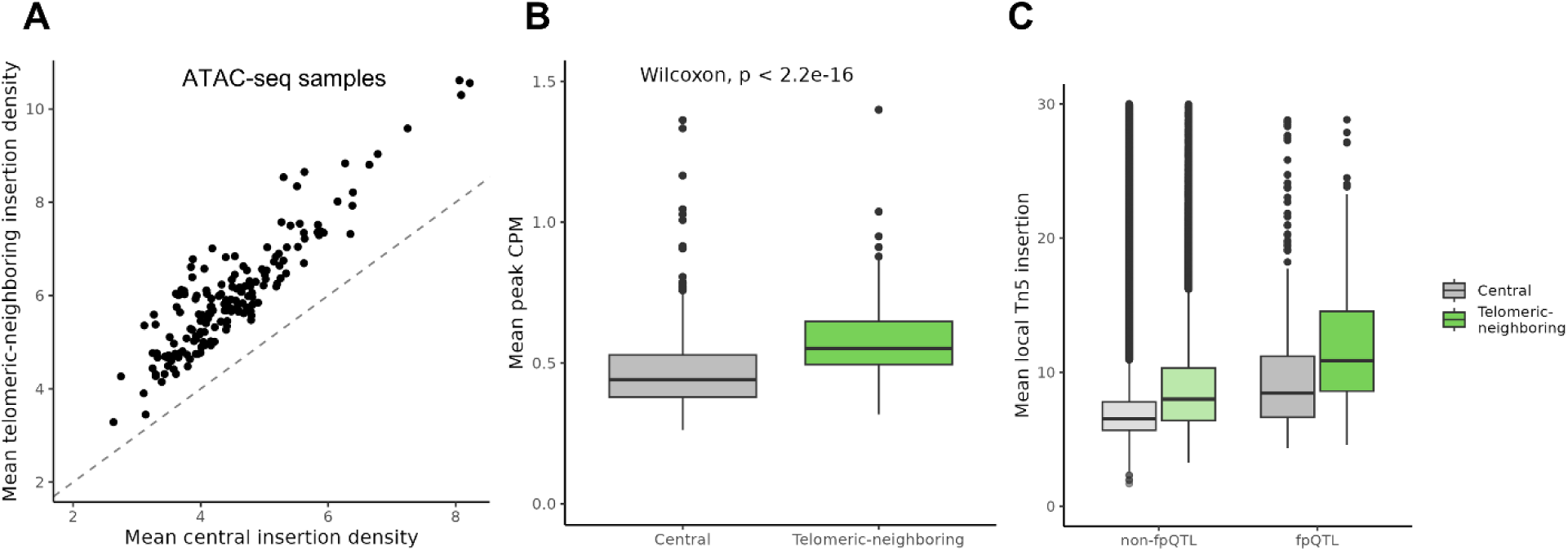
**(A)** The genome was split into bins of 200 bp (the window size PRINT uses to calculate FP score) and the number of insertion sites was measured for each bin. For every sample, the average number of insertions per bin is shown for both near-telomeric bins (within 2 Mb of a chromosome end, y), and central bins (x). **(B)** The average CPM across samples of the peak occupied by a fpQTL, for both near-telomeric fpQTLs (within 2 Mb of a chromosome end, green) and central fpQTLs (grey). **(C)** The average CPM across samples of the peak occupied by any SNP, for both near-telomeric SNPs (within 2 Mb of a chromosome end, green) and central SNPs (grey).

**Figure S8.**
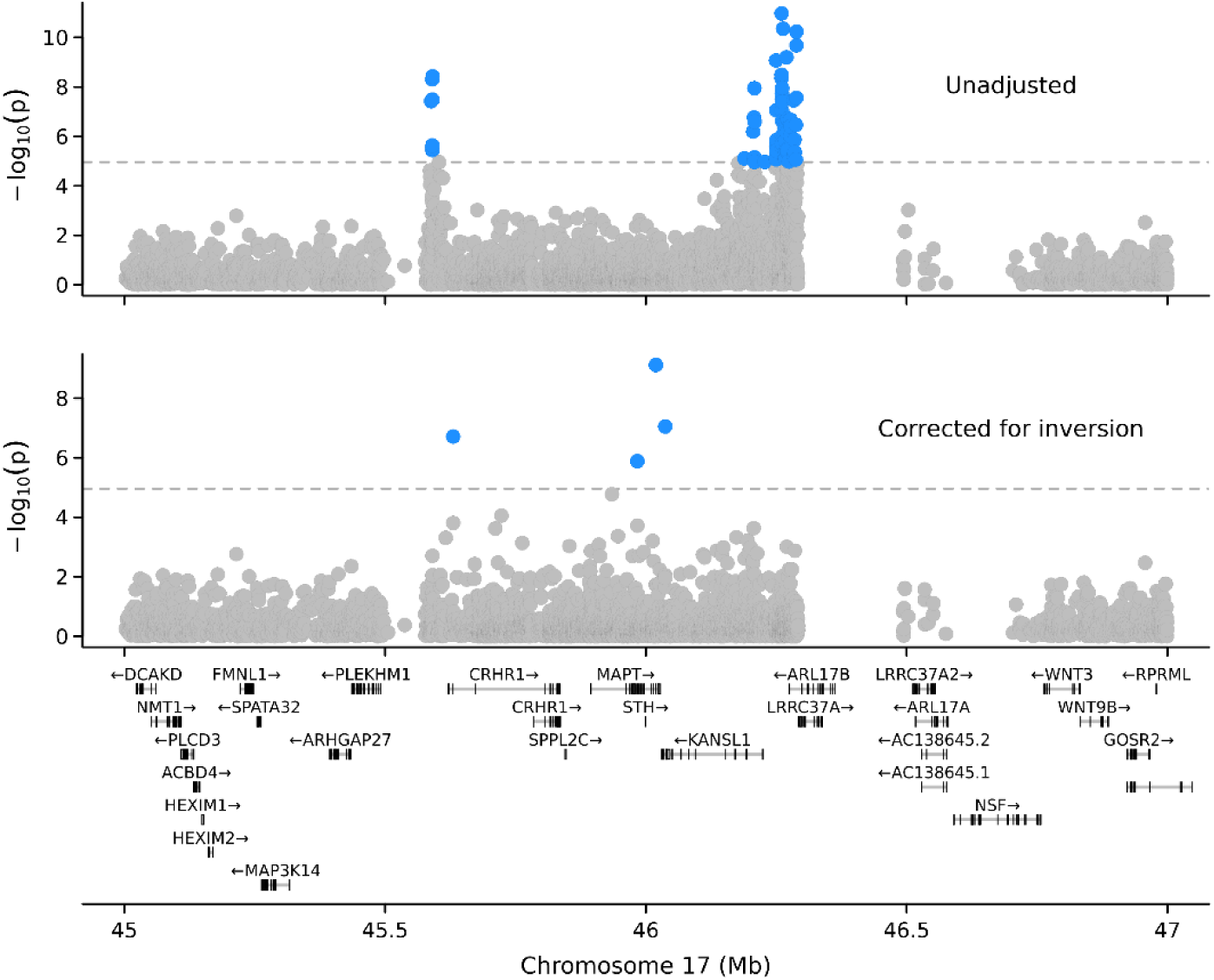
fpQTL significance plot at the chr17 inversion without adjusting for the inversion (top) and after including the inversion genotype as a covariate in the regression (bottom).

## References

1. Maurano, M.T., Humbert, R., Rynes, E., Thurman, R.E., Haugen, E., Wang, H., Reynolds, A.P., Sandstrom, R., Qu, H., Brody, J., et al. (2012). Systematic localization of common disease-associated variation in regulatory DNA. Science 337, 1190–1195. 10.1126/science.1222794.

2. Hormozdiari, F., van de Bunt, M., Segrè, A.V., Li, X., Joo, J.W.J., Bilow, M., Sul, J.H., Sankararaman, S., Pasaniuc, B., and Eskin, E. (2016). Colocalization of GWAS and eQTL Signals Detects Target Genes. The American Journal of Human Genetics 99, 1245–1260. 10.1016/j.ajhg.2016.10.003.

3. Edwards, S.L., Beesley, J., French, J.D., and Dunning, A.M. (2013). Beyond GWASs: Illuminating the Dark Road from Association to Function. Am J Hum Genet 93, 779–797. 10.1016/j.ajhg.2013.10.012.

4. Trynka, G., Sandor, C., Han, B., Xu, H., Stranger, B.E., Liu, X.S., and Raychaudhuri, S. (2013). Chromatin marks identify critical cell types for fine mapping complex trait variants. Nat Genet 45, 124–130. 10.1038/ng.2504.

5. Visscher, P.M., Wray, N.R., Zhang, Q., Sklar, P., McCarthy, M.I., Brown, M.A., and Yang, J. (2017). 10 Years of GWAS Discovery: Biology, Function, and Translation. Am J Hum Genet 101, 5–22. 10.1016/j.ajhg.2017.06.005.

6. Schaid, D.J., Chen, W., and Larson, N.B. (2018). From genome-wide associations to candidate causal variants by statistical fine-mapping. Nat Rev Genet 19, 491–504. 10.1038/s41576-018-0016-z.

7. Shendure, J., Findlay, G.M., and Snyder, M.W. (2019). Genomic Medicine–Progress, Pitfalls, and Promise. Cell 177, 45–57. 10.1016/j.cell.2019.02.003.

8. THE GTEX CONSORTIUM (2020). The GTEx Consortium atlas of genetic regulatory effects across human tissues. Science 369, 1318–1330. 10.1126/science.aaz1776.

9. Yao, D.W., O’Connor, L.J., Price, A.L., and Gusev, A. (2020). Quantifying genetic effects on disease mediated by assayed gene expression levels. Nat Genet 52, 626–633. 10.1038/s41588-020-0625-2.

10. Mostafavi, H., Spence, J.P., Naqvi, S., and Pritchard, J.K. (2023). Systematic differences in discovery of genetic effects on gene expression and complex traits. Nat Genet, 1–10. 10.1038/s41588-023-01529-1.

11. Palermo, J., Chesi, A., Zimmerman, A., Sonti, S., Pahl, M.C., Lasconi, C., Brown, E.B., Pippin, J.A., Wells, A.D., Doldur-Balli, F., et al. (2023). Variant-to-gene mapping followed by cross-species genetic screening identifies GPI-anchor biosynthesis as a regulator of sleep. Science Advances 9, eabq0844. 10.1126/sciadv.abq0844.

12. Su, C., Gao, L., May, C.L., Pippin, J.A., Boehm, K., Lee, M., Liu, C., Pahl, M.C., Golson, M.L., Naji, A., et al. (2022). 3D chromatin maps of the human pancreas reveal lineage-specific regulatory architecture of T2D risk. Cell Metabolism 34, 1394–1409.e4. 10.1016/j.cmet.2022.08.014.

13. Ramdas, S., Judd, J., Graham, S.E., Kanoni, S., Wang, Y., Surakka, I., Wenz, B., Clarke, S.L., Chesi, A., Wells, A., et al. (2022). A multi-layer functional genomic analysis to understand noncoding genetic variation in lipids. The American Journal of Human Genetics 109, 1366–1387. 10.1016/j.ajhg.2022.06.012.

14. Farh, K.K.-H., Marson, A., Zhu, J., Kleinewietfeld, M., Housley, W.J., Beik, S., Shoresh, N., Whitton, H., Ryan, R.J.H., Shishkin, A.A., et al. (2015). Genetic and epigenetic fine mapping of causal autoimmune disease variants. Nature 518, 337–343. 10.1038/nature13835.

15. Sakaue, S., Weinand, K., Isaac, S., Dey, K.K., Jagadeesh, K., Kanai, M., Watts, G.F.M., Zhu, Z., Brenner, M.B., McDavid, A., et al. (2024). Tissue-specific enhancer–gene maps from multimodal single-cell data identify causal disease alleles. Nat Genet, 1–12. 10.1038/s41588-024-01682-1.

16. Buenrostro, J.D., Giresi, P.G., Zaba, L.C., Chang, H.Y., and Greenleaf, W.J. (2013). Transposition of native chromatin for fast and sensitive epigenomic profiling of open chromatin, DNA-binding proteins and nucleosome position. Nat Methods 10, 1213–1218. 10.1038/nmeth.2688.

17. Yan, F., Powell, D.R., Curtis, D.J., and Wong, N.C. (2020). From reads to insight: a hitchhiker’s guide to ATAC-seq data analysis. Genome Biology 21, 22. 10.1186/s13059-020-1929-3.

18. Li, Z., Schulz, M.H., Look, T., Begemann, M., Zenke, M., and Costa, I.G. (2019). Identification of transcription factor binding sites using ATAC-seq. Genome Biology 20, 45. 10.1186/s13059-019-1642-2.

19. Ouyang, N., and Boyle, A.P. (2020). TRACE: transcription factor footprinting using chromatin accessibility data and DNA sequence. Genome Res 30, 1040–1046. 10.1101/gr.258228.119.

20. Bentsen, M., Goymann, P., Schultheis, H., Klee, K., Petrova, A., Wiegandt, R., Fust, A., Preussner, J., Kuenne, C., Braun, T., et al. (2020). ATAC-seq footprinting unravels kinetics of transcription factor binding during zygotic genome activation. Nat Commun 11, 4267. 10.1038/s41467-020-18035-1.

21. Yang, T., and Henao, R. (2022). TAMC: A deep-learning approach to predict motif-centric transcriptional factor binding activity based on ATAC-seq profile. PLOS Computational Biology 18, e1009921. 10.1371/journal.pcbi.1009921.

22. Vierstra, J., Lazar, J., Sandstrom, R., Halow, J., Lee, K., Bates, D., Diegel, M., Dunn, D., Neri, F., Haugen, E., et al. (2020). Global reference mapping of human transcription factor footprints | Nature. Nature 583, 729–736. 10.1038/s41586-020-2528-x.

23. Xu, S., Feng, W., Lu, Z., Yu, C.Y., Shao, W., Nakshatri, H., Reiter, J.L., Gao, H., Chu, X., Wang, Y., et al. (2020). regSNPs-ASB: A Computational Framework for Identifying Allele-Specific Transcription Factor Binding From ATAC-seq Data. Frontiers in Bioengineering and Biotechnology 8.

24. Yan, J., Qiu, Y., Ribeiro dos Santos, A.M., Yin, Y., Li, Y.E., Vinckier, N., Nariai, N., Benaglio, P., Raman, A., Li, X., et al. (2021). Systematic analysis of binding of transcription factors to noncoding variants. Nature 591, 147–151. 10.1038/s41586-021-03211-0.

25. Abramov, S., Boytsov, A., Bykova, D., Penzar, D.D., Yevshin, I., Kolmykov, S.K., Fridman, M.V., Favorov, A.V., Vorontsov, I.E., Baulin, E., et al. (2021). Landscape of allele-specific transcription factor binding in the human genome. Nat Commun 12, 2751. 10.1038/s41467-021-23007-0.

26. Ouyang, N., and Boyle, A.P. (2022). Quantitative assessment of association between noncoding variants and transcription factor binding. Preprint, 10.1101/2022.11.22.517559 10.1101/2022.11.22.517559.

27. Quach, B., and Furey, T.S. (2017). DeFCoM: analysis and modeling of transcription factor binding sites using a motif-centric genomic footprinter. Bioinformatics 33, 956–963. 10.1093/bioinformatics/btw740.

28. Weirauch, M.T., Cote, A., Norel, R., Annala, M., Zhao, Y., Riley, T.R., Saez-Rodriguez, J., Cokelaer, T., Vedenko, A., Talukder, S., et al. (2013). Evaluation of methods for modeling transcription factor sequence specificity. Nat Biotechnol 31, 126–134. 10.1038/nbt.2486.

29. Asrani, S.K., Devarbhavi, H., Eaton, J., and Kamath, P.S. (2019). Burden of liver diseases in the world. Journal of Hepatology 70, 151–171. 10.1016/j.jhep.2018.09.014.

30. Miao, Z., Garske, K.M., Pan, D.Z., Koka, A., Kaminska, D., Männistö, V., Sinsheimer, J.S., Pihlajamäki, J., and Pajukanta, P. (2022). Identification of 90 NAFLD GWAS loci and establishment of NAFLD PRS and causal role of NAFLD in coronary artery disease. Human Genetics and Genomics Advances 3, 100056. 10.1016/j.xhgg.2021.100056.

31. Vujkovic, M., Ramdas, S., Lorenz, K.M., Guo, X., Darlay, R., Cordell, H.J., He, J., Gindin, Y., Chung, C., Myers, R.P., et al. (2022). A multiancestry genome-wide association study of unexplained chronic ALT elevation as a proxy for nonalcoholic fatty liver disease with histological and radiological validation. Nat Genet 54, 761–771. 10.1038/s41588-022-01078-z.

32. Mahajan, A., Spracklen, C.N., Zhang, W., Ng, M.C.Y., Petty, L.E., Kitajima, H., Yu, G.Z., Rüeger, S., Speidel, L., Kim, Y.J., et al. (2022). Multi-ancestry genetic study of type 2 diabetes highlights the power of diverse populations for discovery and translation. Nat Genet 54, 560–572. 10.1038/s41588-022-01058-3.

33. Graham, S.E., Clarke, S.L., Wu, K.-H.H., Kanoni, S., Zajac, G.J.M., Ramdas, S., Surakka, I., Ntalla, I., Vedantam, S., Winkler, T.W., et al. (2021). The power of genetic diversity in genome-wide association studies of lipids. Nature 600, 675–679. 10.1038/s41586-021-04064-3.

34. Pazoki, R., Vujkovic, M., Elliott, J., Evangelou, E., Gill, D., Ghanbari, M., van der Most, P.J., Pinto, R.C., Wielscher, M., Farlik, M., et al. (2021). Genetic analysis in European ancestry individuals identifies 517 loci associated with liver enzymes. Nat Commun 12, 2579. 10.1038/s41467-021-22338-2.

35. Loos, R.J.F., and Yeo, G.S.H. (2022). The genetics of obesity: from discovery to biology. Nat Rev Genet 23, 120–133. 10.1038/s41576-021-00414-z.

36. Yengo, L., Sidorenko, J., Kemper, K.E., Zheng, Z., Wood, A.R., Weedon, M.N., Frayling, T.M., Hirschhorn, J., Yang, J., Visscher, P.M., et al. (2018). Meta-analysis of genome-wide association studies for height and body mass index in ∼700000 individuals of European ancestry. Human Molecular Genetics 27, 3641–3649. 10.1093/hmg/ddy271.

37. Littleton, S.H., Berkowitz, R.I., and Grant, S.F.A. (2020). Genetic Determinants of Childhood Obesity. Mol Diagn Ther 24, 653–663. 10.1007/s40291-020-00496-1.

38. Çalışkan, M., Manduchi, E., Rao, H.S., Segert, J.A., Beltrame, M.H., Trizzino, M., Park, Y., Baker, S.W., Chesi, A., Johnson, M.E., et al. (2019). Genetic and Epigenetic Fine Mapping of Complex Trait Associated Loci in the Human Liver. The American Journal of Human Genetics 105, 89–107. 10.1016/j.ajhg.2019.05.010.

39. Corces, M.R., Trevino, A.E., Hamilton, E.G., Greenside, P.G., Sinnott-Armstrong, N.A., Vesuna, S., Satpathy, A.T., Rubin, A.J., Montine, K.S., Wu, B., et al. (2017). An improved ATAC-seq protocol reduces background and enables interrogation of frozen tissues. Nat Methods 14, 959–962. 10.1038/nmeth.4396.

40. Loh, P.-R., Danecek, P., Palamara, P.F., Fuchsberger, C., Reshef, Y.A., Finucane, H.K., Schoenherr, S., Forer, L., McCarthy, S., Abecasis, G.R., et al. (2016). Reference-based phasing using the Haplotype Reference Consortium panel. Nat Genet 48, 1443–1448. 10.1038/ng.3679.

41. Hu, Y., Ma, S., Kartha, V.K., Duarte, F.M., Horlbeck, M., Zhang, R., Shrestha, R., Labade, A., Kletzien, H., Meliki, A., et al. (2023). Single-cell multi-scale footprinting reveals the modular organization of DNA regulatory elements. Preprint, 10.1101/2023.03.28.533945 10.1101/2023.03.28.533945.

42. Aguet, F., Brown, A.A., Castel, S.E., Davis, J.R., He, Y., Jo, B., Mohammadi, P., Park, Y., Parsana, P., Segrè, A.V., et al. (2017). Genetic effects on gene expression across human tissues. Nature 550, 204–213. 10.1038/nature24277.

43. Storey, J.D., and Tibshirani, R. (2003). Statistical significance for genomewide studies. Proceedings of the National Academy of Sciences of the United States of America 100, 9440. 10.1073/pnas.1530509100.

44. Storey, J.D., Bass, A.J., Dabney, A., and Robinson, D. (2023). qvalue: Q-value estimation for false discovery rate control. Bioconductor. http://bioconductor.org/packages/qvalue/.

45. O’Leary, N.A., Wright, M.W., Brister, J.R., Ciufo, S., Haddad, D., McVeigh, R., Rajput, B., Robbertse, B., Smith-White, B., Ako-Adjei, D., et al. (2016). Reference sequence (RefSeq) database at NCBI: current status, taxonomic expansion, and functional annotation. Nucleic Acids Res 44, D733–D745. 10.1093/nar/gkv1189.

46. Arnold, M., Raffler, J., Pfeufer, A., Suhre, K., and Kastenmüller, G. (2015). SNiPA: an interactive, genetic variant-centered annotation browser. Bioinformatics 31, 1334–1336. 10.1093/bioinformatics/btu779.

47. Moore, J.E., Purcaro, M.J., Pratt, H.E., Epstein, C.B., Shoresh, N., Adrian, J., Kawli, T., Davis, C.A., Dobin, A., Kaul, R., et al. (2020). Expanded encyclopaedias of DNA elements in the human and mouse genomes. Nature 583, 699–710. 10.1038/s41586-020-2493-4.

48. Rauluseviciute, I., Riudavets-Puig, R., Blanc-Mathieu, R., Castro-Mondragon, J.A., Ferenc, K., Kumar, V., Lemma, R.B., Lucas, J., Chèneby, J., Baranasic, D., et al. (2024). JASPAR 2024: 20th anniversary of the open-access database of transcription factor binding profiles. Nucleic Acids Research 52, D174–D182. 10.1093/nar/gkad1059.

49. Schep, A. motifmatchr: Fast Motif Matching in R. Bioconductor. http://bioconductor.org/packages/motifmatchr/.

50. Korhonen, J., Martinmäki, P., Pizzi, C., Rastas, P., and Ukkonen, E. (2009). MOODS: fast search for position weight matrix matches in DNA sequences. Bioinformatics 25, 3181–3182. 10.1093/bioinformatics/btp554.

51. Boytsov, A., Abramov, S., Aiusheeva, A.Z., Kasianova, A.M., Baulin, E., Kuznetsov, I.A., Aulchenko, Y.S., Kolmykov, S., Yevshin, I., Kolpakov, F., et al. (2022). ANANASTRA: annotation and enrichment analysis of allele-specific transcription factor binding at SNPs. Nucleic Acids Res 50, W51–W56. 10.1093/nar/gkac262.

52. Ruiz-Arenas, C., Cáceres, A., López-Sánchez, M., Tolosana, I., Pérez-Jurado, L., and González, J.R. (2019). scoreInvHap: Inversion genotyping for genome-wide association studies. PLOS Genetics 15, e1008203. 10.1371/journal.pgen.1008203.

53. Klemm, S.L., Shipony, Z., and Greenleaf, W.J. (2019). Chromatin accessibility and the regulatory epigenome. Nat Rev Genet 20, 207–220. 10.1038/s41576-018-0089-8.

54. Furey, T.S. (2012). ChIP–seq and beyond: new and improved methodologies to detect and characterize protein–DNA interactions. Nat Rev Genet 13, 840–852. 10.1038/nrg3306.

55. Kyrmizi, I., Hatzis, P., Katrakili, N., Tronche, F., Gonzalez, F.J., and Talianidis, I. (2006). Plasticity and expanding complexity of the hepatic transcription factor network during liver development. Genes Dev 20, 2293–2305. 10.1101/gad.390906.

56. Lee, C.S., Friedman, J.R., Fulmer, J.T., and Kaestner, K.H. (2005). The initiation of liver development is dependent on Foxa transcription factors. Nature 435, 944–947. 10.1038/nature03649.

57. Ebrahimkhani, M.R., Oakley, F., Murphy, L.B., Mann, J., Moles, A., Perugorria, M.J., Ellis, E., Lakey, A.F., Burt, A.D., Douglass, A., et al. (2011). Stimulating healthy tissue regeneration by targeting the 5-HT2B receptor in chronic liver disease. Nat Med 17, 1668–1673. 10.1038/nm.2490.

58. Finucane, H.K., Bulik-Sullivan, B., Gusev, A., Trynka, G., Reshef, Y., Loh, P.-R., Anttila, V., Xu, H., Zang, C., Farh, K., et al. (2015). Partitioning heritability by functional annotation using genome-wide association summary statistics. Nat Genet 47, 1228–1235. 10.1038/ng.3404.

59. Meuleman, W., Muratov, A., Rynes, E., Halow, J., Lee, K., Bates, D., Diegel, M., Dunn, D., Neri, F., Teodosiadis, A., et al. (2020). Index and biological spectrum of human DNase I hypersensitive sites. Nature 584, 244–251. 10.1038/s41586-020-2559-3.

60. Zhang, K., Hocker, J.D., Miller, M., Hou, X., Chiou, J., Poirion, O.B., Qiu, Y., Li, Y.E., Gaulton, K.J., Wang, A., et al. (2021). A single-cell atlas of chromatin accessibility in the human genome. Cell 184, 5985–6001.e19. 10.1016/j.cell.2021.10.024.

61. Stefansson, H., Helgason, A., Thorleifsson, G., Steinthorsdottir, V., Masson, G., Barnard, J., Baker, A., Jonasdottir, A., Ingason, A., Gudnadottir, V.G., et al. (2005). A common inversion under selection in Europeans. Nat Genet 37, 129–137. 10.1038/ng1508.

62. González, J.R., Ruiz-Arenas, C., Cáceres, A., Morán, I., López-Sánchez, M., Alonso, L., Tolosana, I., Guindo-Martínez, M., Mercader, J.M., Esko, T., et al. (2020). Polymorphic Inversions Underlie the Shared Genetic Susceptibility of Obesity-Related Diseases. The American Journal of Human Genetics 106, 846–858. 10.1016/j.ajhg.2020.04.017.

63. Wang, H., Makowski, C., Zhang, Y., Qi, A., Kaufmann, T., Smeland, O.B., Fiecas, M., Yang, J., Visscher, P.M., and Chen, C.-H. (2023). Chromosomal inversion polymorphisms shape human brain morphology. Cell Reports 42, 112896. 10.1016/j.celrep.2023.112896.

64. Okbay, A., Baselmans, B.M.L., De Neve, J.-E., Turley, P., Nivard, M.G., Fontana, M.A., Meddens, S.F.W., Linnér, R.K., Rietveld, C.A., Derringer, J., et al. (2016). Genetic variants associated with subjective well-being, depressive symptoms, and neuroticism identified through genome-wide analyses. Nat Genet 48, 624–633. 10.1038/ng.3552.

65. Musunuru, K., Strong, A., Frank-Kamenetsky, M., Lee, N.E., Ahfeldt, T., Sachs, K.V., Li, X., Li, H., Kuperwasser, N., Ruda, V.M., et al. (2010). From noncoding variant to phenotype via SORT1 at the 1p13 cholesterol locus. Nature 466, 714–719. 10.1038/nature09266.

66. Wang, X., Raghavan, A., Peters, D.T., Pashos, E.E., Rader, D.J., and Musunuru, K. (2018). Interrogation of the Atherosclerosis-Associated SORT1 (Sortilin 1) Locus With Primary Human Hepatocytes, Induced Pluripotent Stem Cell-Hepatocytes, and Locus-Humanized Mice. Arteriosclerosis, Thrombosis, and Vascular Biology 38, 76–82. 10.1161/ATVBAHA.117.310103.

67. Argemi, J., Latasa, M.U., Atkinson, S.R., Blokhin, I.O., Massey, V., Gue, J.P., Cabezas, J., Lozano, J.J., Van Booven, D., Bell, A., et al. (2019). Defective HNF4alpha-dependent gene expression as a driver of hepatocellular failure in alcoholic hepatitis. Nat Commun 10, 3126. 10.1038/s41467-019-11004-3.

68. Ehle, C., Iyer-Bierhoff, A., Wu, Y., Xing, S., Kiehntopf, M., Mosig, A.S., Godmann, M., and Heinzel, T. (2024). Downregulation of HNF4A enables transcriptomic reprogramming during the hepatic acute-phase response. Commun Biol 7, 1–14. 10.1038/s42003-024-06288-1.

69. Odom, D.T., Zizlsperger, N., Gordon, D.B., Bell, G.W., Rinaldi, N.J., Murray, H.L., Volkert, T.L., Schreiber, J., Rolfe, P.A., Gifford, D.K., et al. (2004). Control of pancreas and liver gene expression by HNF transcription factors. Science 303, 1378–1381. 10.1126/science.1089769.

70. Tan, W.X., Sim, X., Khoo, C.M., and Teo, A.K.K. (2023). Prioritization of genes associated with type 2 diabetes mellitus for functional studies. Nat Rev Endocrinol 19, 477–486. 10.1038/s41574-023-00836-1.

71. Hansel, M.C., Gramignoli, R., Skvorak, K.J., Dorko, K., Marongiu, F., Blake, W., Davila, J., and Strom, S.C. (2014). The History and Use of Human Hepatocytes for the Study and Treatment of Liver Metabolic Diseases. Curr Protoc Toxicol 62, 14.12.1-14.12.23. 10.1002/0471140856.tx1412s62.

72. Adey, A., Morrison, H.G., Asan, Xun, X., Kitzman, J.O., Turner, E.H., Stackhouse, B., MacKenzie, A.P., Caruccio, N.C., Zhang, X., et al. (2010). Rapid, low-input, low-bias construction of shotgun fragment libraries by high-density in vitro transposition. Genome Biology 11, R119. 10.1186/gb-2010-11-12-r119.

